# Regulation of *NTRK2* alternative splicing by PRPF40B controls neural differentiation and synaptic plasticity

**DOI:** 10.1101/2025.04.07.647526

**Authors:** María Duarte-Ruiz, Adela Moreno-Castillo, Younes El Yousfi, Cristina Moreno-Castro, Noelia Martínez-Martínez, Sandra Jiménez-Lozano, Marion Kennel, Candela Ruiz-Rodríguez, Alonso Rodríguez-Caparrós, Jennifer López-Ros, Pierre de la Grange, Cristina Hernández-Munain, Carlos Suñé

## Abstract

BDNF signaling through its receptor TRKB plays a critical role in brain development, neuroplasticity and homeostasis. Alternative splicing of the TRKB gene, *NTRK2*, generates either the full-length receptor (TRKB-FL) or a truncated isoform (TRKB-T1) that inhibits BDNF signaling and has been implicated in neurodegenerative diseases, psychiatric disorders and cognitive impairments. Here, we show that PRPF40B, a splicing factor associated with neuronal dysfunction, promotes production of the TRKB-FL isoform during neuronal differentiation. Silencing PRPF40B increases TRKB-T1 expression, impairing expression of genes important for neuronal differentiation and synaptic plasticity. Our data thus identify PRPF40B as a key regulator of the balance between TRKB receptor isoforms, crucial for fine-tuning neuronal responses and for preventing neuroplasticity or survival impairments, providing also a mechanism for the role of PRPF40B in the pathogenesis of various human neurodegenerative diseases and psychiatric disorders

## INTRODUCTION

The regulation of neuronal differentiation is essential for the development and regeneration of the nervous system. Neurotrophins, such as brain-derived neurotrophic factor (BDNF), play a key role in modulating the formation of new synapses and the development of neural plasticity. The primary high-affinity receptor for BDNF is TRKB (Tropomyosin receptor kinase B), a member of the Trk family of receptor tyrosine kinases^1^. Upon BDNF binding, TRKB undergoes dimerization and autophosphorylation on its intracellular tyrosine residues, activating downstream signaling pathways, including PLCγ, MAPK/ERK, and PI3K/AKT to promote neuronal differentiation and synaptic plasticity^2^.

The human TRKB gene (*NTRK2*) is relatively large, comprising 24 exons and capable of generating more than 30 potential isoforms through alternative splicing^3,4^. Despite this diversity, the most highly expressed isoforms in the mammalian brain are the full-length receptor, TRKB-FL, and the truncated variant TRKB-T1^3^. In the TRKB-T1 isoform, the splicing machinery excludes the exons encoding the tyrosine kinase domain. Instead, an alternative exon (exon 16) is included, introducing a premature stop codon. This results in a truncated receptor lacking kinase activity. The TRKB-T1 variant acts as a dominant-negative form of TRKB-FL, unable to activate the signaling pathways required for neurite outgrowth and synaptogenesis^5–7^. The functional role of TRKB-T1 variant is to regulate the intensity of neurotrophic responses elicited by BDNF binding. Thus, maintaining the balance between TRKB-FL and TRKB-T1 is essential for proper neuronal differentiation.

The dysregulation of BDNF-TRKB signaling through aberrant upregulation of the TRKB-T1 isoform has been implicated in various human diseases and dysfunctions, particularly neurological and psychiatric disorders, including Alzheimer’s disease (AD), Parkinson’s disease, Amyotrophic Lateral Sclerosis (ALS) and Frontotemporal dementia (FTD), Huntington’s disease (HD), epilepsy, stroke, mood disorders and schizophrenia^8^. Findings showing that restoring TRKB-T1 to physiological levels rescues neuronal sensitivity to BDNF and prevents neuronal cell death *in vivo*^9,10^ validate therapeutic strategies aimed at normalizing TRKB-T1 levels. An attractive strategy to correct the aberrant TRKB-T1 upregulation is to target *NTRK2* alternative splicing. However, very little is currently known about the mechanisms that regulate *NTRK2* expression or the factors driving its pathological dysregulation. To date, the only known RNA-binding protein (RBP) implicated in *NTRK2* regulation is Rbfox1. Rbfox1 binds directly to *NTRK2* RNA and promotes TRKB-T1 stability rather than influencing splicing. Notably, while increased Rbfox1 expression enhances TRKB-T1 levels, reducing Rbfox1 does not have an effect^11^. Therefore, the mechanisms controlling the alternative splicing that generates TRKB-T1 versus TRKB-FL, as well as the factors involved, remain unknown.

PRPF40B (PRP40 pre-mRNA processing factor 40 homolog B) is one of the mammalian orthologs of the essential yeast splicing factor Prp40. The yeast Prp40 participates in the early steps of spliceosome formation of precursor mRNA^12^, a role that PRPF40B might have conserved^13^. PRPF40B contains two WW and five FF domains in its primary sequence. It interacts with the branchpoint binding protein/SF1 through its WW-containing amino-terminal region and with U2A^F65^ through its FF domains, supporting its role in stabilizing splicing factors near the 3’ splice site^13^. This initial splicing reaction is essential for facilitating subsequent interactions among snRNPs and splicing factors, ultimately leading to the catalytic step of splicing. This has been supported by a study demonstrating that PRPF40B regulates hundreds of alternative splicing events in K562 lymphoblast cells, predominantly involving cassette exons flanked by weak 5’ and 3’ splice sites^14^. The other Prp40 homolog, PRPF40A, has been more extensively studied than PRPF40B and, based on phylogenetic data, appears to be more closely related to Prp40, with PRPF40B emerging significantly later in evolutionary history^15^. Like PRPF40B, PRPF40A interacts with SF1 and U2A^F65^ ^16^ and has been identified in the human A and B spliceosomal complexes^17^ as well as in the U2 snRNP^18^. Recent studies have shown that PRPF40A promotes the inclusion of short cassette exons in mouse neuroblastoma (NB) cells^19^, consistent with previous findings where Prp40 was required for the splicing of microexons surrounded by conventionally sized introns^20^. Furthermore, PRPF40A modulates the inclusion of short cassette exons in human HL-60 leukemia cells. In this case, the exons are flanked by short GC-rich introns^21^.

PRPF40B has been implicated in the pathogenesis of various human neurodegenerative diseases and psychiatric disorders, including HD, AD, ALS/FTD, Rett syndrome, and schizophrenia^22–28^, suggesting an important role for PRPF40B in neuronal development and function. Supporting this notion, PRPF40B expression is higher in the nervous system compared to other tissues^22,23^. Moreover, a recent proteomic analysis of mouse cell type-specific brain nuclei identified PRPF40B specifically in neurons^29^, where TRKB expression is also high. Here, we report that PRPF40B regulates the transcription and splicing of numerous neuronal genes involved in proliferation, cell growth, neurogenesis, and synaptogenesis. Our findings reveal that PRPF40B depletion disrupts BDNF-mediated signaling by upregulating TRKB-T1 receptor levels through the inclusion of exon 16 in the pre-mRNA, leading to impaired neuronal differentiation and synaptic plasticity.

## RESULTS

### Silencing of PRPF40B inhibits proliferation, colony formation, and induces cell-cycle arrest in human NB cells *in vitro*

NB cells originate from neural crest cells and possess the capacity for unlimited proliferation *in vitro* while maintaining the potential to differentiate into neuronal cell types. In this context, the SH-SY5Y human NB cell line is extensively utilized to investigate the mechanisms underlying neuronal differentiation and neurite outgrowth, as well as to serve as a model for studying neuronal disorders^30^. To clarify the regulatory effects of PRPF40B in the nervous system, we studied three SH-SY5Y NB cell lines: WT (non-genetically modified) and two PRPF40B-silenced lines, G1 and G2, generated using specific CRISPR/Cas9 guides. PRPF40B expression in G1 was intermediate between WT and G2, with the latter exhibiting the highest level of silencing (Fig. 1a, inset). We first examined whether PRPF40B knockdown affected cell proliferation using the resazurin viability assay over five consecutive days. Both G1 and G2 cell lines showed reduced proliferation compared with the WT control (Fig. 1a). Notably, the G1 cell line showed an intermediate level of proliferation between G2 and WT cells, which is in accordance to their level of PRPF40B expression. Immunostaining for the proliferative marker Ki-67 was performed in the SH-SY5Y cell lines to confirm the reduced proliferative ability of PRPF40B-silenced cells. The Ki-67 immunostaining data indicated that silencing PRPF40B led to a significant decrease in cell proliferation compared to the control cells (Fig. 1b). The ability of colony formation of SH-SY5Y cells was also decreased in the G2 cells, which exhibit the highest level of PRPF40B silencing (Fig.1c). These findings are consistent with the previous results and indicate that downregulation of PRPF40B inhibits the proliferative ability of SH-SY5Y cells.

**Fig. 1.**
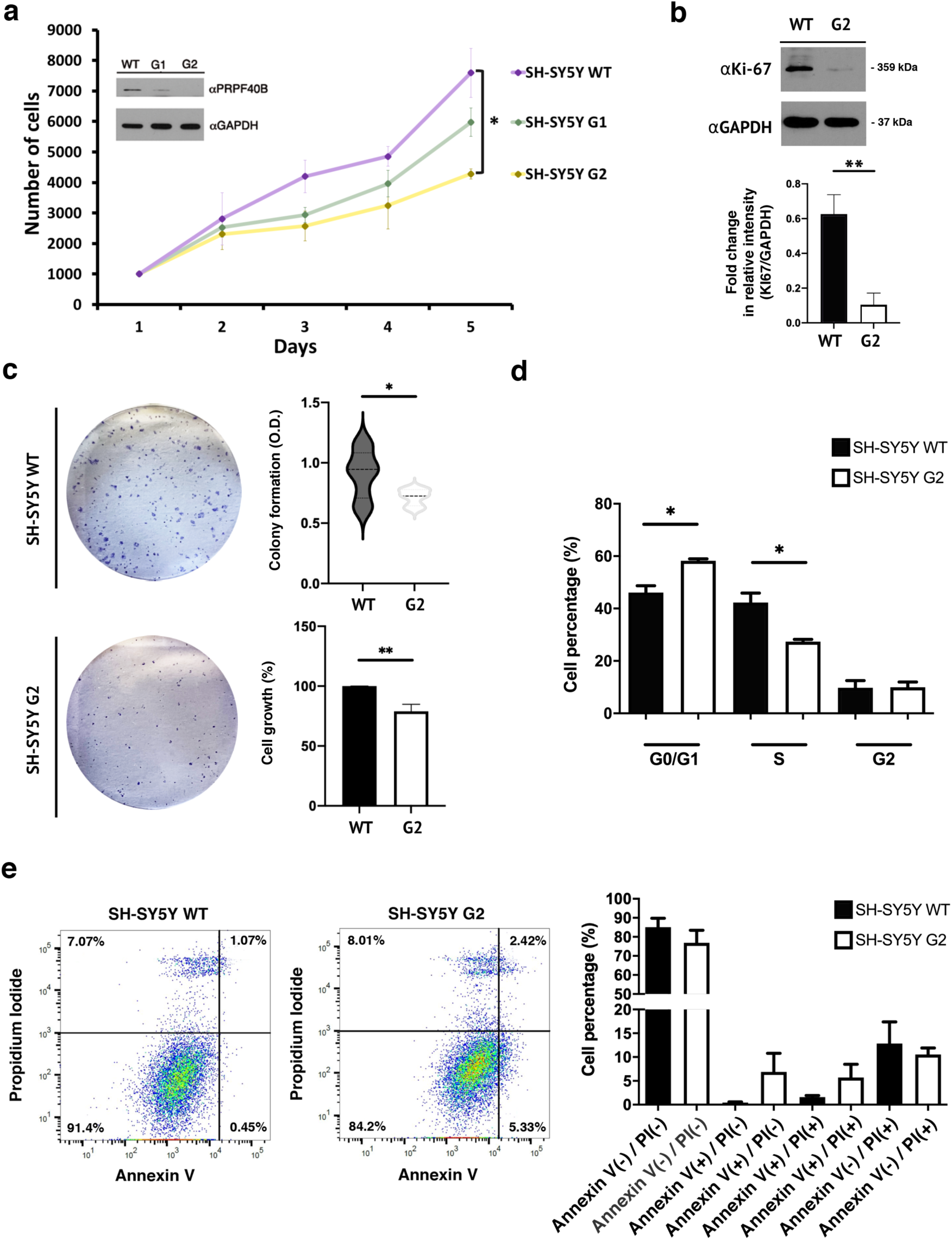
PRPF40B downregulation impairs proliferation, colony formation, and induces cell-cycle arrest in SH-SY5Y cells. **a** The resazurin assay was used to measure cell viability of SH-SY5Y WT, G1, and G2 cells over five consecutive days. Inset: PRPF40B expression was assessed by Western blot analysis using specific antibodies. **b** Western blot (top) and quantification (bottom) analysis of Ki-67 expression in SH-SY5Y WT and G2 cells. **c** Colony formation assay. Representative images of colonies formed by SH-SY5Y WT and G2 cells after 7 days (left). The graphs (right) show relative differences in colony formation and cell growth in G2 compared to control cells. **d** Cell cycle distribution of SH-SY5Y WT and G2 cells. The graph shows the percentage of cells in each phase of the cell cycle. **e** SH-SY5Y WT and G2 cells were analyzed by Annexin V-FITC/PI staining with flow cytometry. The lower-right quadrant represents early apoptotic cells, while the upper quadrants represent late apoptotic cells. Summary graphs of the flow cytometry results are shown at the bottom. Data represent the mean ± SEM from three independent experiments. Statistical significance: *p ≤ 0.05 and **p ≤ 0.01.

To investigate the mechanisms underlying the altered proliferation in PRPF40B-silenced cells, we performed propidium iodide incorporation experiments to examine the regulatory effect of PRPF40B on the cell cycle, using the G2 cell line with the highest level of PRPF40B silencing. Cell cycle analysis revealed that silencing PRPF40B resulted in a G0/G1 phase arrest and a reduction in the S phase in G2 cells (Fig.1d) compared to the WT cells. We also evaluated the rate of apoptosis to investigate the effects of PRPF40B silencing on cell death in SH-SY5Y cells. Notably, PRPF40B silencing did not significantly affect the apoptotic rate (Fig. 1e). These results suggest that the inhibition of cell proliferation in modified SH-SY5Y cells may be attributed to G0/G1 phase arrest caused by PRPF40B silencing, without affecting apoptosis in NB cells.

To further corroborate these data, PRPF40B was depleted in SH-SY5Y cells using specific shRNAs, and changes in proliferation between control (Shc) and PRPF40B-depleted cells (Sh7y(e)) were analyzed by the resazurin assay. Confirming the previous results, the Sh7y(e) cells showed decreased proliferation compared to the shRNA control cell line (Supplementary Fig. 1a). The protein levels of PRPF40B in the shRNA cell lines were examined by Western blotting (Supplementary Fig. 1a, inset). Decreased levels for the proliferative marker Ki-67 was observed in Sh7y(e) cells, confirming the reduced proliferative ability of these PRPF40B-silenced cells (Supplementary Fig. 1b).

To assess whether overexpression of PRPF40B could rescue the proliferative impairment in PRPF40B-silenced cells, we transduced G2 cells with either an empty MigR1 retrovirus or retroviruses overexpressing PRPF40B (G2-GFP-Control and G2-GFP-PRPF40B, respectively). Flow cytometry was used to monitor the expression of GFP-expressing MigR1 (Supplementary Fig. 2a), while Western blot and RT-qPCR analyses were employed to detect PRPF40B expression (Supplementary Fig. 2b and 2c). We then monitored cell proliferation in the transduced cells. Overexpression of PRPF40B in the G2 PRPF40B-silenced cells rescued Ki-67 expression levels (Supplementary Fig. 2d) and enhanced their proliferative ability in the resazurin assay (Supplementary Fig. 2e). These results confirm our previous data and strongly suggest a direct role for PRPF40B in cell proliferation of SH-SY5Y cells.

### PRPF40B silencing affects NB cell migration and differentiation *in vitro*

Neuronal migration is a critical process in nervous system development. During neurogenesis, communication between neurons and their environment is essential for guiding them to their proper destination in the brain, enabling them to perform their functions^31^. Using the scratch wound-healing assay, a widely employed method to assess cell migration in culture, we observed reduced migration in G2 cells (Fig. 2a), suggesting that PRPF40B plays a role in regulating cell migration in the SH-SY5Y cell line.

**Fig. 2.**
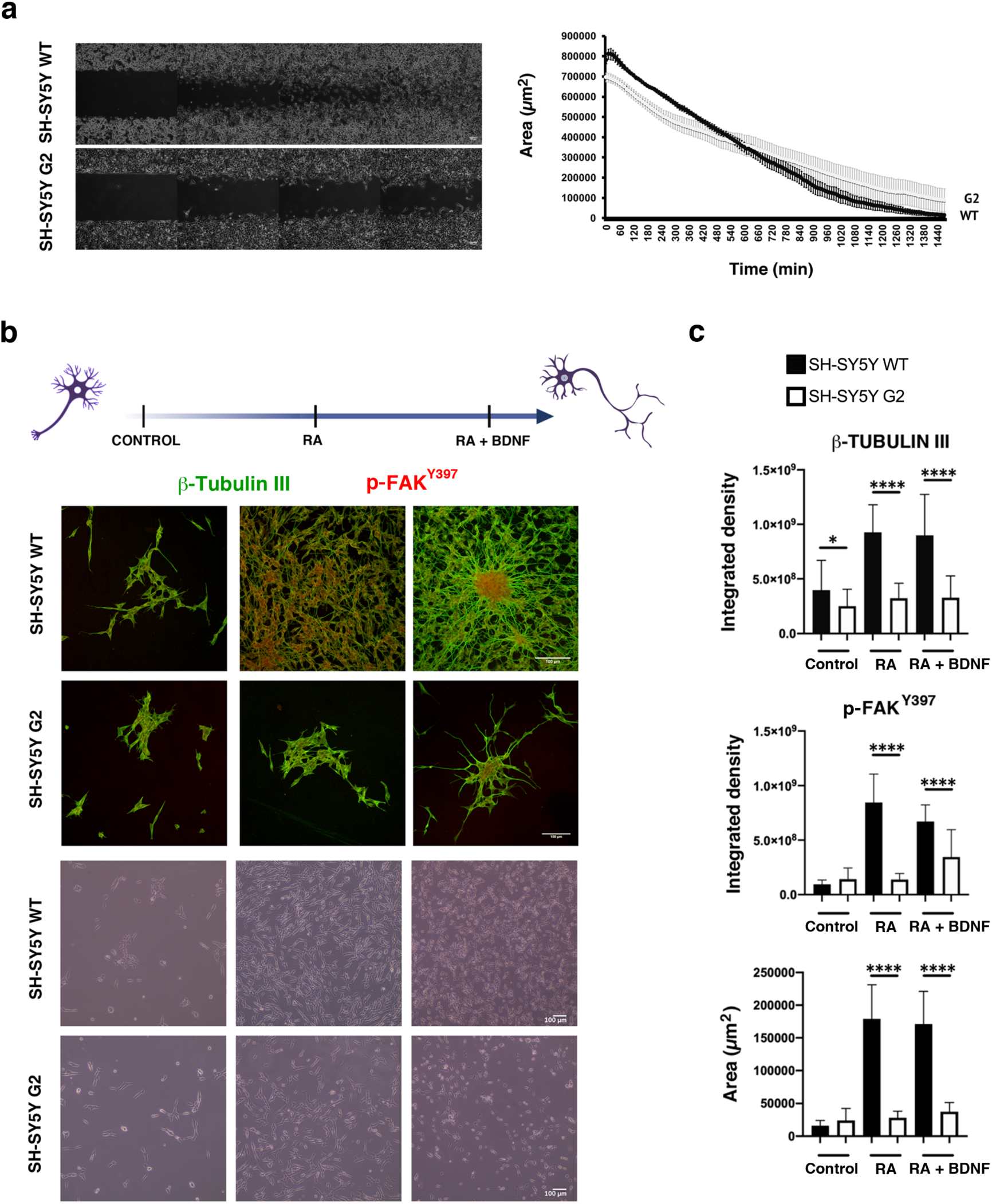
PRPF40B depletion impairs cell migration and neuronal differentiation in NB cells. **a** Representative images and graphical representation of wound-healing assays performed using SH-SY5Y WT and G2 cells. Data represent the mean ± SEM from three independent experiments. **b** The neuronal differentiation assay was performed using RA and RA + BDNF. Top panels: Immunofluorescence staining with dual labeling of cells was performed using antibodies directed against β-Tubulin III (green) and p-FAK^Y397^ (red). β-Tubulin III indicates neuronal plasticity, while p-FAK^Y397^ indicates migration, adhesion, and neuronal synaptogenesis. Bottom panels: Phase-contrast microscopy photographs of the culture plate during differentiation. Scale bars: 100 µm. **c** Graphical representation of the fluorescence intensity for each label, with the area indicating neuronal growth. Data represent the mean ± SEM from two independent experiments, each with n = 10 images. Statistical significance: *\**p ≤ 0.05, and ****p **≤** 0.0001.

There is a tightly regulated balance between proliferation and differentiation in the nervous system, where neural progenitors first expand through proliferation and then specialize into functional neurons or glial cells, ensuring proper brain development and function. The effect of PRPF40B on cell proliferation suggests a potential role for this factor in neuronal activity, which is essential for shaping synaptic strength and connectivity. To examine the effect of PRPF40B silencing on neural cell differentiation, we induced the differentiation of SH-SY5Y WT and G2 cells using a sequential retinoic acid (RA) differentiation and brain-derived neurotrophic factor (BDNF) maturation protocol. SH-SY5Y cells were immunostained with the specific neuronal marker β-tubulin III antibody and with p-FAK^Y397^. The focal adhesion kinase FAK is the most highly expressed kinase in the developing central nervous system and plays important roles in neuronal migration, axonal guidance, neurite outgrowth, proliferation, survival, and differentiation^32^. FAK binds to growth factor receptors, initiating its activation by autophosphorylation at residue Y397 (p-FAK^Y397^), which promotes focal contact turnover and cell motility^33,34^. Despite the proliferation block in G2 cells, the RA/BDNF treatment stimulated dendritic growth in both cell populations (WT and G2), with p-FAK^Y397^ localized at the cell center but polarized toward the leading protrusion (Fig. 2b). The fluorescence intensity of β-tubulin III and p-FAK^Y397^ clusters relative to cell area clearly indicates that PRPF40B silencing impairs SH-SY5Y cell differentiation (Fig. 2c). Additionally, quantitative analysis of cell area in each condition confirmed a growth delay in PRPF40B-silenced cells (Fig. 2c).

### PRPF40B regulates gene transcription in NB cells

To gain insights into the regulatory effects of PRPF40B in the nervous system and the molecular mechanisms by which its silencing affects proliferation and differentiation, we first identified the overall changes in the transcriptome profile of SH-SY5Y cells resulting from PRPF40B silencing. We studied three SH-SY5Y cell lines: WT, scramble (SCR; transfected with CRISPR/Cas9 machinery using control guides), and G2. The levels of PRPF40B expression in these cells were assessed by Western blot analysis (Fig. 3a). Based on hierarchical clustering of regulated genes, numerous differentially expressed (DE) genes were identified, particularly in comparison to G2 (Fig. 3b). From the analysis of DE genes among G2/WT, G2/SCR, and SCR/WT cells, we selected 2,243 genes specifically affected by PRPF40B silencing (Fig. 3c and Supplementary Table 1). Of these, 739 (32,9%) were down-regulated and 1,504 (67,1%) were up-regulated upon PRPF40B knockdown compared to control cells (Fig. 3c), suggesting that PRPF40B preferentially functions as a transcriptional repressor in SH-SY5Y cells. We validated 10 randomly selected DE genes by RT-qPCR in G2 cells, achieving a validation rate of 100%, which supports the robustness of our approach (Fig.3d). To further validate the transcriptomic results, we performed RT-qPCR for three of the DE genes using both the G1 and G2 cell lines. We obtained an intermediate level of gene expression with G1 cells, as expected due to their intermediate level of PRPF40B expression (Supplementary Fig. 3a). In addition, G2-GFP-PRPF40B transduced cells were also able to partially restore the mRNA expression levels of many endogenous targets of PRPF40B (Supplementary Fig. 3b), corroborating the specificity of the observed effects linked to PRPF40B silencing.

**Fig. 3.**
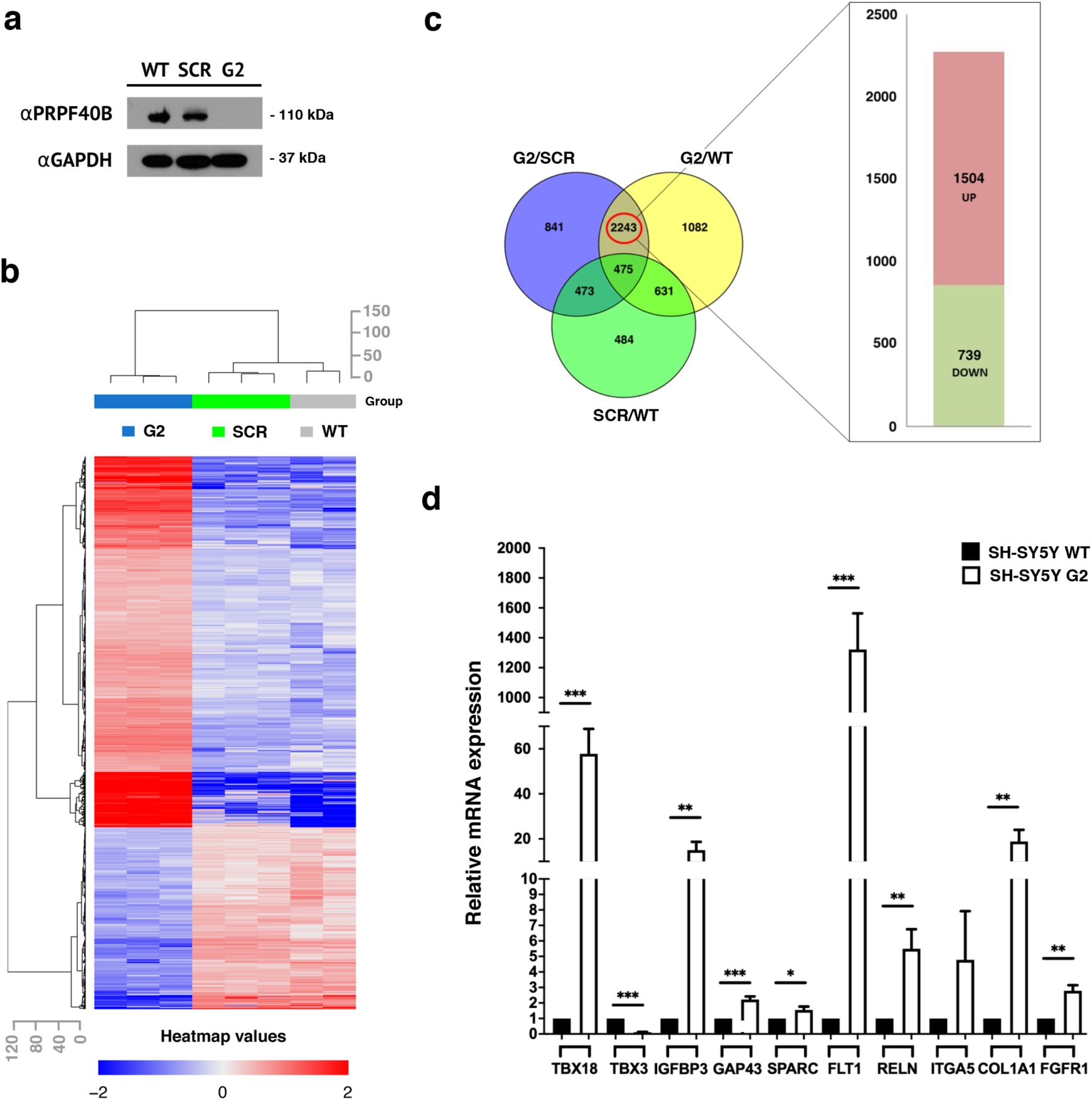
Identification of differentially expressed genes after PRPF40B silencing in NB cells. **a** PRPF40B expression in SH-SY5Y WT, scrambled (SCR), and G2 cells. Cell lysates were resolved by SDS-PAGE, and proteins were transferred to membranes for the detection of PRPF40B and GAPDH expression using specific antibodies. **b** Heat map representation of differentially expressed (DE) genes in WT, SCR, and G2 cells. The top row shows a dendrogram representing clustering information and similarity relationships among the cell lines. The left column of the heat map displays the gene clusters. The color intensity of the rows represents changes in gene expression: red indicates upregulated genes, and blue indicates downregulated genes. **c** The Venn diagram illustrates the overlap among all DE genes. A total of 2,243 DE genes were identified, including 1,504 upregulated genes (red) and 739 downregulated genes (green). **d** RNA-seq validation was performed using RT-qPCR for selected DE genes of interest. Data represent the mean ± SEM from three to six independent experiments. Statistical significance: *p ≤ 0.05, **p ≤ 0.01, and ***p ≤ 0.001.

*In silico* analysis of the 2,243 DE genes was carried out using various computational platforms. Enrichment analysis of the down-regulated gene cluster, utilizing the GO biological process pathway, Reactome, and KEGG databases, identified DE genes involved in the release of neurotransmitters from synaptic vesicles (Supplementary Fig. 4a), a crucial process for proper communication between neurons. Disruption of synaptic vesicle dynamics is believed to contribute to cognitive impairments in neurodegenerative disorders^35^. Enrichment analysis of the up-regulated gene cluster indicated that signal transduction pathways related to cell survival and growth in response to extracellular signals were significantly affected. Notably, the regulation of cell migration and extracellular matrix organization were pathways highlighted in the GO biological process database, supporting the potential role of PRPF40B in the migration of NB cells (Supplementary Fig. 4b).

### PRPF40B regulates alternative RNA splicing in NB cells

PRPF40B has been previously characterized as a splicing factor^13^, prompting us to analyze the effects of PRPF40B knockdown on RNA splicing. Analysis of the RNA-seq transcriptome data revealed 233 alternative splicing (AS) events across 200 distinct genes from comparisons among DE events among G2/WT, G2/SCR, and SCR/WT cells (Fig.4a). These events were classified into the following patterns: 90 exon cassette, 64 alternative first exons, 17 alternative terminal exons, 6 alternative acceptor splice sites, 5 alternative donor splice sites, 4 mutually exclusive exons, and 4 intron retention (Fig. 4b and Supplementary Table 2). Among the 233 AS events influenced by PRPF40B expression, exon cassettes (38,6%) and alternative first exons (27,5%) were the most prevalent splicing patterns (Fig. 4b). Analysis of PRPF40B-regulated AS events annotated in the reference FAST DB database indicated that the frequency of cassette exons and alternative first exons significantly exceed what would be expected based on their representation in the genome (Fig. 4c).

**Fig. 4.**
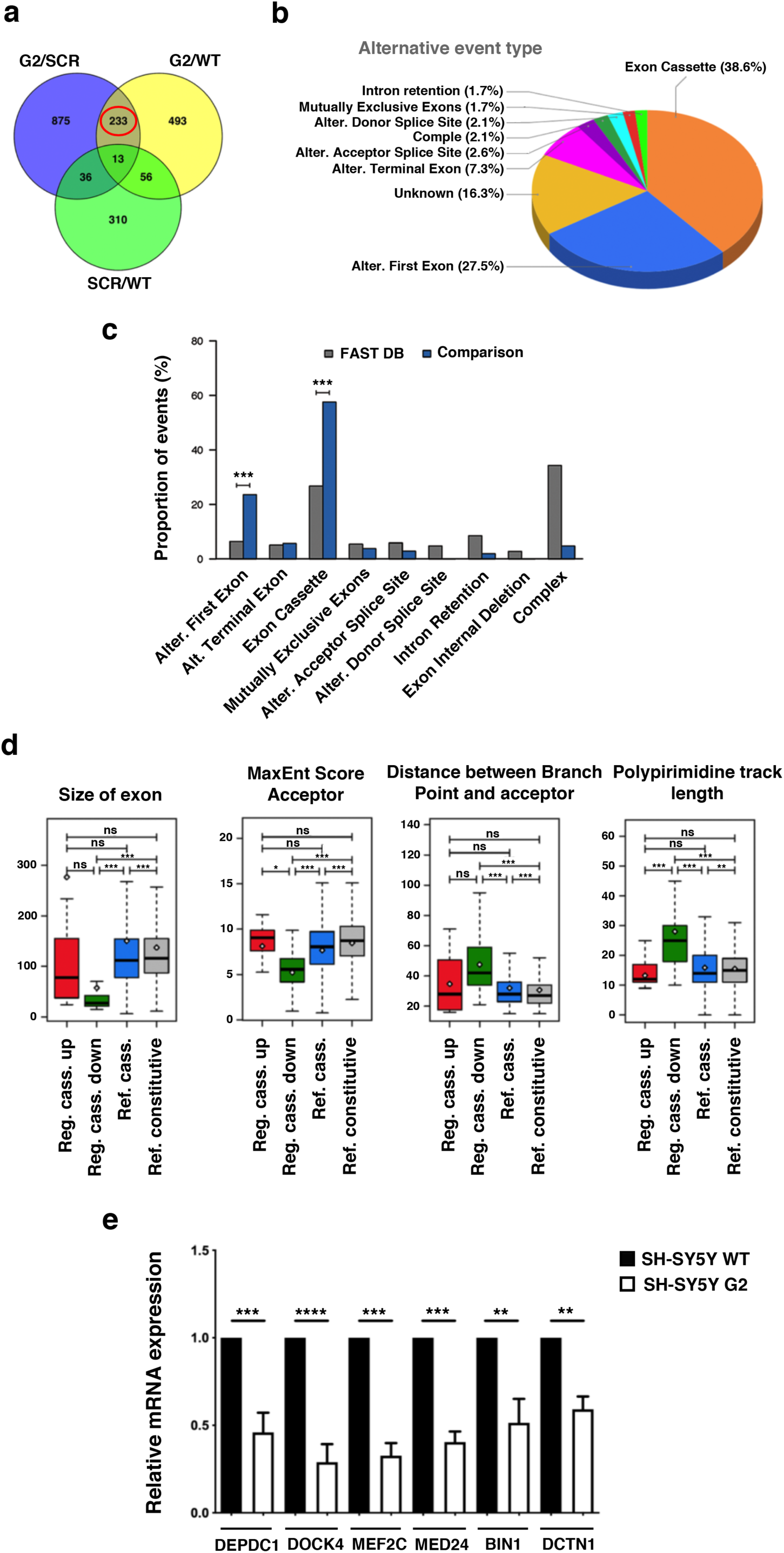
PRPF40B silencing alters alternative splicing patterns in NB cells. **a** Venn diagram showing the overlap of alternative splicing (AS) events among G2/SCR, G2/WT, and SCR/WT comparisons. **b** Pie chart illustrating the types of AS events among the 233 DE events. **c** Proportion of PRPF40B-regulated AS events compared to annotated events in the reference FAST-DB database. **d** Structural and sequence characteristics of downregulated exons, including exon size, strength of acceptor sites, distance between branch points and acceptor sites, and length of polypyrimidine tracts. **e** Validation of six selected alternative exon splicing events by RT-qPCR. The y-axis represents the expression of the transcript in G2 cells relative to that in WT cells, normalized to 1. Data represent the mean ± SEM from three to six independent experiments. Statistical significance: **p ≤ 0.01, ***p ≤ 0.001, and ****p ≤ 0.0001.

Given these results, particularly the lower number of alternative first exons, we focused on the exon cassette pattern and performed analyses to identify specific features of these cassette exons. PRPF40B-regulated exons displayed distinct changes compared to reference non-regulated exons. Specifically, down-regulated exons were significantly smaller than either reference or up-regulated exons, had weaker acceptor sites, and exhibited longer distances between branch points (BPs) and acceptor sites due to extended polypyrimidine tracts (Fig. 4d). No significant motifs were enriched in the PRPF40B-regulated exons and flanking introns compared to both non-regulated cassette exons and constitutive exons. To validate these findings, we performed RT-qPCR analysis of several representative down-regulated exon events, confirming the splicing changes identified by RNA-seq in WT and G2 cells (Fig. 4e and Supplementary Fig. 5a). In addition, we further validated these results by analyzing the expression levels of two down-regulated exon events in the G2-GFP-Control and G2-GFP-PRPF40B transduced cells. The expression of PRPF40B in the G2 cells restored the expression of the regulated exon (Supplementary Fig. 5b). *In silico* analysis of the 200 genes with DE splicing events identified many interconnected cellular processes, including Rho GTPase signaling, microtubule dynamics, focal adhesion regulation, and microtubule cytoskeleton organization, all of which play crucial roles in neuronal development and differentiation. These processes enable neurons to undergo the morphological and functional changes necessary for establishing neuronal networks during development (Supplementary Fig. 6a). Finally, comparative bioinformatic analysis revealed a limited but significant overlap between the genes showing changes in both transcript and exon levels (Supplementary Fig. 6b and Supplementary Table 3).

### PRPF40B regulates critical signaling pathways involved in neuronal differentiation

Given our data, we sought to investigate the role of PRPF40B in neuronal differentiation more comprehensively. During this process, RA and BDNF act synergistically to enhance neuronal development by modulating overlapping signaling pathways and transcriptional programs. RA initiates transcriptional programs that prime cells for differentiation, while BDNF amplifies these effects by activating critical signaling pathways, including MAPK/ERK and PI3K/AKT (Fig. 5a). Interestingly, these pathways were prominent targets in PRPF40B-silenced cells (Supplementary Fig. 4a). To explore the effects of PRPF40B during differentiation, we analyzed the levels of several components of the MAPK/ERK, and PI3K/AKT pathways (ERK 1/2, p-ERK, AKT, p-AKT, and PI3K). As expected, these kinases signaling pathways, along with their downstream targets related to neuronal differentiation, synaptic function, and cytoskeletal remodeling were activated during differentiation. However, the absence of PRPF40B significantly reduced the levels of MAPK/ERK and PI3K/AKT pathway components, as well as their downstream targets (Fig. 5b and 5f). Furthermore, we analyzed the levels of several factors involved in synaptic machinery, neurotransmitter release, and cytoskeletal dynamics (N-cadherin, complexin 1/2, synaptophysin, synapsin 1a/b, β-tubulin III, and drebrin), which support BDNF signaling through TRKB (Fig. 5c-d and 5g-h). These findings highlight the essential role of PRPF40B in activating these signaling pathways, which are crucial for proper neuronal differentiation, synaptogenesis, and plasticity. If PRPF40B plays an important role during neuronal differentiation, its expression level could be higher upon differentiation. To determine whether PRPF40B expression is upregulated during neuronal differentiation, we analyzed its expression levels during the differentiation process. As shown in Fig. 5e and 5i, the PRPF40B expression significantly increased in WT cells during differentiation.

**Fig. 5.**
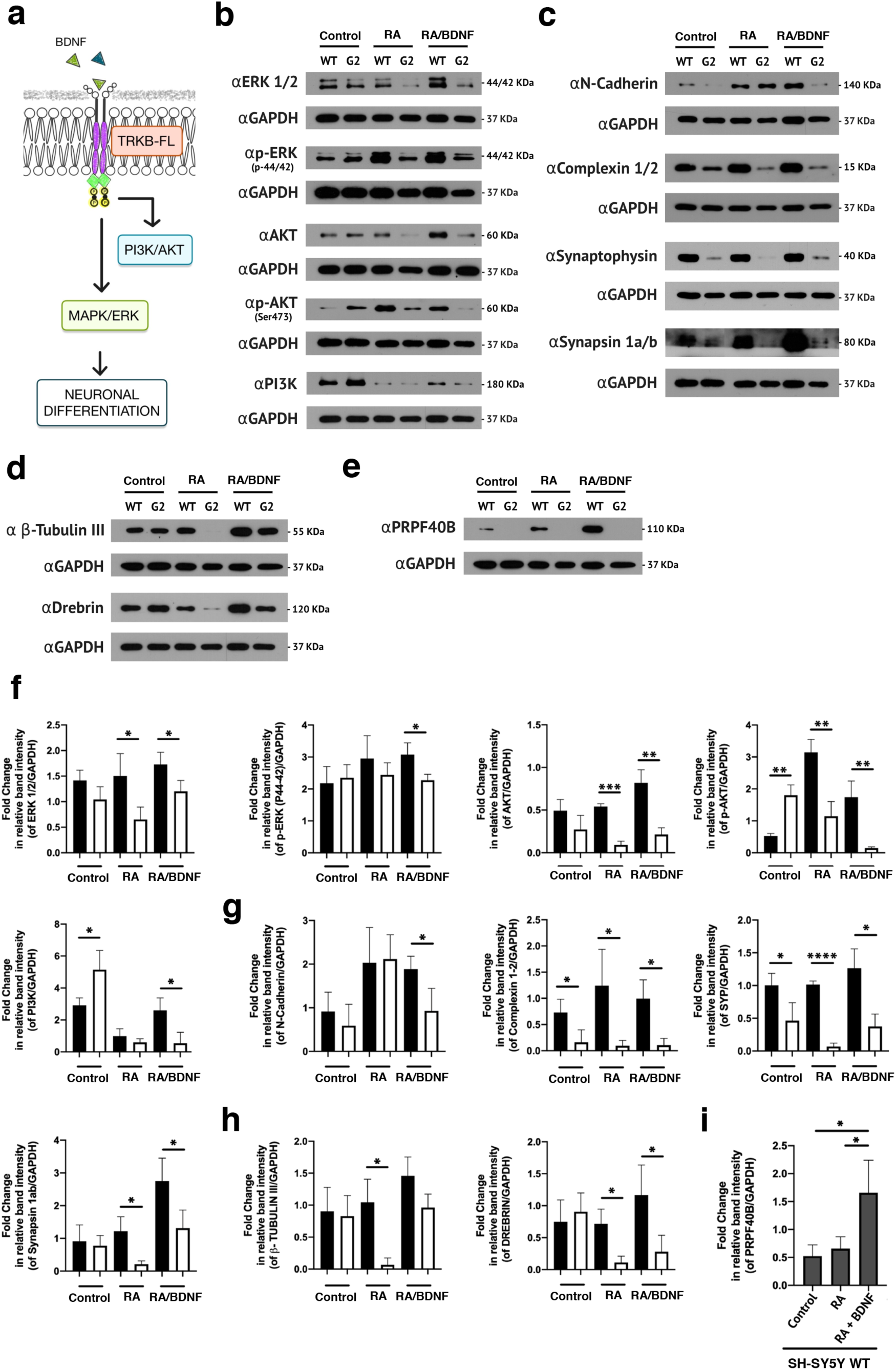
PRPF40B regulates key signaling pathways involved in neuronal differentiation and synaptic plasticity. **a** Schematic representation of the BDNF signaling pathway. **b** and **f** Western blot analysis and quantification of MAPK and PI3K pathway components (ERK1/2, p-ERK, AKT, p-AKT, and PI3K) in control, RA, and RA/BDNF-treated WT and G2 cell lines. **c** and **g** Western blot analysis and quantification of synaptogenesis-related proteins (N-cadherin, complexin 1/2, synaptophysin, and synapsin 1a/b) in control, RA, and RA/BDNF-treated WT and G2 cell lines. **d** and **h** Western blot analysis and quantification of proteins related to neuronal size and plasticity (β-Tubulin III and drebrin) in control, RA, and RA/BDNF-treated WT and G2 cell lines. **e** and **i** Western blot analysis and quantification of PRPF40B expression in control, RA, and RA/BDNF-treated WT and G2 cell lines. Data represent the mean ± SEM from three independent experiments. Statistical significance: *p ≤ 0.05, **p ≤ 0.01, ***p ≤ 0.001, and ****p ≤ 0.0001. In the quantification graphs, black and white bars represent SH-SY5Y WT and G2 cells, respectively.

### PRPF40B regulates the expression of the TRKB-T1 deletion variant

Decreased levels of MAPK/ERK, PI3K/AKT, and their downstream targets upon differentiation may indicate a problem with the TRKB receptor in G2 cells. During differentiation, a balance between TRKB-FL and TRKB-T1 is essential, as overexpression of TRKB-T1 is known to negatively regulate the BDNF-TRKB pathway (Fig. 6a). Analysis of TRKB-FL and its dominant-negative TRKB-T1 receptor expression levels in WT and G2 cell lines revealed increased protein expression of TRKB-FL and TRKB-T1 in WT cells upon RA treatment (Fig. 6b), priming the cells to respond to BDNF. This observation aligns with previous findings at the mRNA level^36^. In marked contrast, the TRKB-FL receptor was undetectable in G2 cells, while the TRKB-T1 variant exhibited a pronounced increase in expression (Fig. 6b). Following BDNF treatment, the expression levels of both TRKB-FL and TRKB-T1 decreased in G2 cells (Fig. 6b), possibly indicating increased lysosomal degradation of the receptors in these cells^37,38^. TRKB-T1 appeared as a double band in gels, likely reflecting reported post-translational modifications, particularly glycosylation^39,40^.

**Fig. 6.**
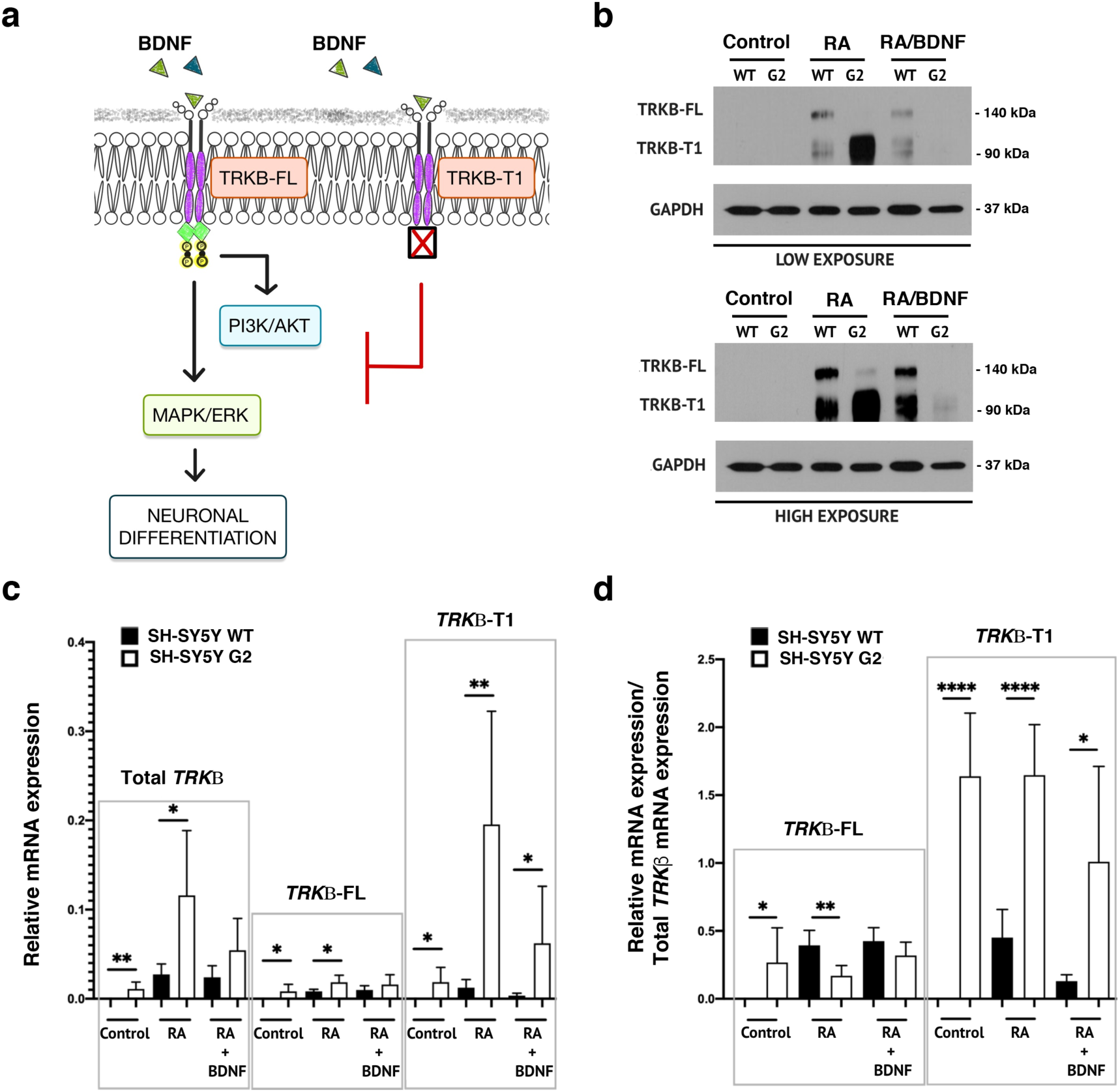
PRPF40B modulates *NTRK2* splicing to regulate TRKB-FL expression during differentiation. **a** Schematic representation of BDNF signaling through TRKB-FL and TRKB-T1. **b** Western blot analysis of TRKB-FL and TRKB-T1 expression levels in control, RA, and RA/BDNF-treated WT and G2 cell lines. **c** Relative mRNA expression levels of total *TRKβ*, *TRKβ-FL*, and *TRKβ-T1* in control, RA, and RA/BDNF-treated WT and G2 cell lines. **d** Relative mRNA expression levels of *TRKβ-FL* and *TRKβ-T1*, normalized to total *TRKβ*, in control, RA, and RA/BDNF-treated WT and G2 cell lines. Data represent the mean ± SEM from three to six independent experiments. Statistical significance: *p ≤ 0.05, **p ≤ 0.01, and ****p ≤ 0.0001.

Further analysis of *TRKB* receptor isoform mRNA levels (Supplementary Fig. 7a) revealed a global increase in total *TRKB* expression upon differentiation, with significantly higher levels observed in G2 compared to WT cells (Fig. 6c, Total TRKB). Notably, this increase was driven by a shift in alternative splicing that strongly favored expression of the truncated *TRKB-T1* isoform in G2 cells (Fig. 6c, *TRKB-T1*). To account for the effects of PRPF40B silencing on *NTRK2* processing, we analyzed the relative expression of each isoform normalized to total *TRKB* mRNA levels using specific primers (Supplementary Fig. 7a). This analysis showed that the elevated *TRKB* expression observed in the G2 cell line during differentiation results specifically from an increased proportion of the truncated *TRKB-T1* isoform (Fig. 6d, *TRKB-T1*). In contrast, in WT cells, the balance between *TRKB-FL* and *TRKB-T1* remains stable throughout neuronal differentiation (Fig. 6d). Detailed bioinformatics analysis using EASANA revealed exon 16 inclusion in the TRKB gene (*NTRK2*) in G2 cells, introducing a stop codon and generating a truncated isoform lacking the kinase domain (Supplementary Fig. 7b). These findings strongly support a direct role for PRPF40B in regulating *TRKB* precursor mRNA splicing, leading to increased expression of a dominant-negative truncated TRKB.

## DISCUSSION

PRPF40B has been associated with various neuronal dysfunctions, including neurodegenerative and psychiatric disorders. These conditions often share disruptions in differentiation, neurogenesis, and synaptogenesis, which are critical processes for maintaining brain structure and function. Here, we show that PRPF40B regulates key signaling pathways, including MAPK/ERK and PI3K/AKT. PRPF40B silencing leads to a significant reduction in the level of these pathway components and their downstream targets upon neuronal differentiation. Additionally, the downregulation of synaptic proteins (e.g., N-cadherin, complexin 1/2, synaptophysin, synapsin 1a/b) and cytoskeletal regulators (β-tubulin III, drebrin) suggests that PRPF40B is critical for maintaining synaptic integrity and neuronal connectivity. Given that both MAPK/ERK and PI3K/AKT pathways are implicated in developmental and neurodegenerative disorders, our data suggest that PRPF40B dysfunction could be a contributing factor to these pathologies.

Dysregulated BDNF signaling is a hallmark feature across many of these disorders, with most therapeutic strategies focusing on restoring natural BDNF levels. However, the efficacy of BDNF-based treatments is limited due its inability to cross the blood-brain barrier (BBB), short half-life, and associated side effects. Furthermore, this therapeutic approach becomes ineffective when alterations in TRKB receptor expression occur, particularly the upregulation of the TRKB-T1 isoform, which acts as a dominant-negative receptor and inhibits normal BDNF signaling.

Previous to our work, no factors have been shown to regulate *NTRK2* splicing. Here, we show that PRPF40B influences the neuronal expression level of TRKB receptors. PRPF40B silencing upregulates the expression of the dominant-negative TRKB-T1 receptor through alternative splicing, and this regulation impairs BDNF-mediated cellular signaling, leading to significant physiological consequences. Therefore, PRPF40B is an essential factor to maintain normal balance between the levels of the different isoforms of the TRKB receptor (Fig. 7).

**Fig. 7.**
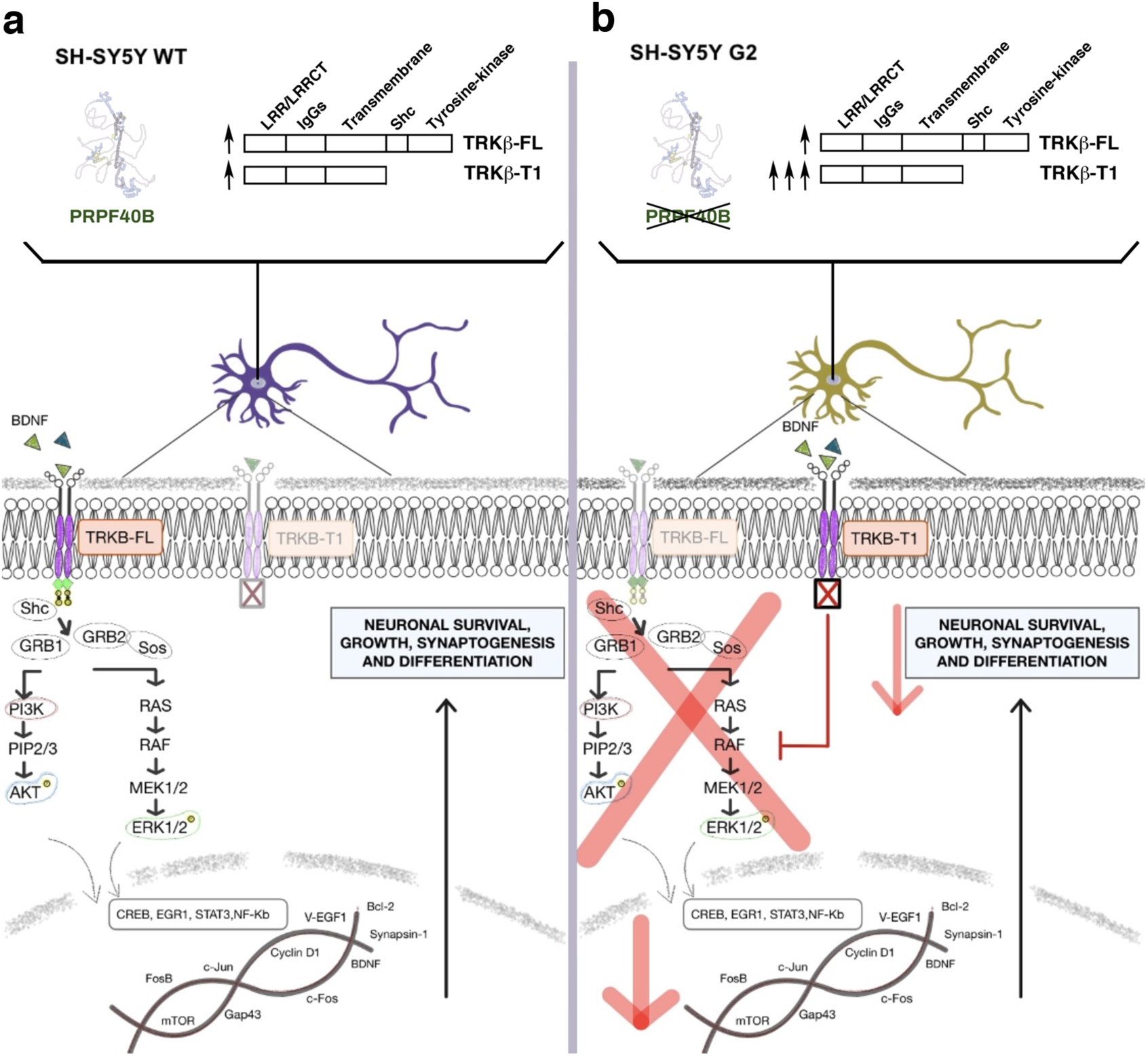
A Hypothetical model for the role of PRPF40B in BDNF-induced neuronal differentiation. **a** In cells with normal PRPF40B expression (SH-SY5Y WT), BDNF interacts with the TRKB-FL receptor, triggering a cascade of signaling events through the MAPK/ERK and PI3K/AKT pathways to activate key genes essential for proper neuronal differentiation, synaptogenesis, and plasticity. **b** In cells where PRPF40B expression is silenced (SH-SY5Y G2), TRKB-T1 receptor expression is upregulated during differentiation. BDNF binds to TRKB-T1 without activating the classical signaling cascades due to the absence of the kinase domain, thereby reducing BDNF availability for TRKB-FL and impairing differentiation and synaptic maturation.

The precise molecular mechanism by which PRPF40B regulates TRKB alternative splicing is unknown and requires further investigation. The main alternative splicing events regulated by PRPF40B involve cassette exons. PRPF40B-downregulated exons were significantly smaller than reference or up-regulated exons, had weaker acceptor sites, and exhibited longer distances between BPs and acceptor sites due to extended polypyrimidine tracts. These longer distances may reduce splicing efficiency, as reflected in the higher prevalence of exon skipping and lower inclusion levels observed for exons preceded by more distant BPs^41^. These findings partially align with previous results in K562 cells^14^ and support the role of PRPF40B in stabilizing splicing complexes near the 3’ splice site, thereby facilitating interactions with the 5’ splice site, particularly for exons with weak splice signals.

In humans, exon 16 is long (5,165 nt), and its inclusion introduces a premature stop codon, resulting in a 33-nucleotide sequence that encodes a unique intracellular 11-amino-acid tail, which is highly conserved among humans, rodents, and chickens^8^. Here, we show that PRPF40B depletion increases exon 16 inclusion, indicating that PRPF40B normally restricts its inclusion. One possible explanation is that PRPF40B enhances recognition of the 3’ splice site flanking exon 17 (which is 48-nucleotides long and has a weak 3’ splice site signal, according to ESEfinder), facilitating exon 15-exon 17 pairing, promoting exon 17 inclusion and maintaining a proper balance between isoforms. When PRPF40B is lost, this pairing is disrupted, leading to more frequent inclusion of exon 16. Alternatively, PRPF40B may actively repress exon 16 inclusion, suggesting a dual role in splicing regulation: defining exon/intron boundaries to ensure proper splicing and modulating exon 16 inclusion/exclusion to maintain the correct downstream splicing outcome. Further research is needed to elucidate the precise mechanism by which PRPF40B regulates exon 16 splicing in *NTRK2*.

TRKB-FL transcripts predominate during early developmental stages, while TRKB-T1 transcripts become more abundant in later stages and in the adult organism^39^. This temporal expression pattern suggests that the critical functions involved in the initial formation and organization of the brain during early development are performed through signaling via the TRKB-FL receptors. In adult stages, where is critical to maintain and control the established neuronal connections, the TRKB-T1 isoform modulates the intensity of BDNF signaling through the TRKB-FL receptor to prevent a damaging overstimulation of BDNF-dependent signaling pathways. Our results indicate that SH-SY5Y WT cells express very low levels of TRKB. However, upon differentiation, both TRKB-FL and TRKB-T1 are induced to similar levels, as detected by Western blot (Fig. 6b). PRPF40B depletion in these cells strongly increases TRKB-T1 isoform levels. Although PRPF40B expression increases during neuronal differentiation, TRKB-T1 remains abundant. This apparent contradiction in our SH-SY5Y cells results suggests that additional splicing regulators or alternative transcriptional and posttranscriptional processes contribute to TRKB-T1 expression.

Our work also shows that while PRPF40B regulates 200 genes at the splicing level, it unexpectedly influences the transcription of over 2,000 genes in SH-SY5Y cells. Furthermore, the limited overlap between genes affected at both the transcriptional and splicing levels suggests that PRP40B operates through distinct molecular mechanisms to regulate these processes. The fact that PRPF40B silencing induces differential expression of hundreds of genes, impacting key cellular processes such as proliferation, viability, migration, and neural differentiation, aligns with emerging evidence that RBP and splicing factors play broader roles in the regulation of gene expression beyond their traditional functions in RNA processing and metabolism. Seminal research from the Fu laboratory demonstrated that the SR protein SRSF2 stimulates transcriptional elongation by releasing the RNA polymerase complex from promoter-proximal pausing^42,43^. The small nuclear ribonucleoprotein particle U1 (U1 snRNP) has a critical splicing-independent function in preventing premature cleavage and polyadenylation from cryptic polyadenylation signals. Similarly, there is growing evidence showing that splicing can modulate epigenetic patterns of chromatin. For example, U1 snRNP alters chromatin organization, and several splicing factors, such as SRSF1, SRSF2 and hnRNP A1, regulate nucleosome organization^44^. These and other observations^45^ highlight an emerging role for splicing factors in transcriptional control and chromatin structure. Additional data support this alternative mechanism, such as the reported association of the WW and FF domains of Prp40 with the phosphorylated C-terminal domain of RNA polymerase II^46^ ^15^. In addition, recent findings with the *Drosophila* Prp40 (dPrp40) indicate that dPrp40 regulates histone mRNA expression by modulating transcription rather than its mRNA processing^47^. In this work, we observed an overall increase in *NTRK2* transcripts levels (Fig. 6c), alongside a switch in isoform expression in PRPF40B-depleted SH-SY5Y cells (Fig. 6d). This raises the question of whether, in addition to its role in splicing regulation, PRPF40B influences also *NTRK2* transcription. PRPF40B might exert either a direct transcriptional regulatory function or act indirectly through its splicing activities on other transcriptional regulators. The phenotypic effects of PRPF40B depletion may thus be a consequence of the increase in the total *NTRK2* transcription and an increase in *TRKB-T1* isoform levels. This could indicate that the function of PRPF40B extends beyond maintaining TRKB isoform balance to also regulating overall *NTRK2* expression. Overall, these findings suggest a potential new function for Prp40 proteins in transcriptional regulation that remains to be investigated, reinforcing the idea that splicing factors play a role in various aspects of regulated gene expression, extending beyond their conventional function in RNA processing.

Although proliferation and differentiation are often considered antagonistic processes, where proliferating cells remain undifferentiated and differentiated cells exit the cell cycle, PRPF40B depletion inhibits both. This apparent contradiction suggests that PRPF40B may regulate a shared upstream pathway essential for both processes, possibly through its role in splicing regulation affecting key signaling pathways. Although TRKB-controlled processes are required for both proliferation and differentiation, TRKB protein expression remained very low or undetectable in both WT and PRPF40B-depleted undifferentiated NB cells (Fig. 6b), with its expression being largely dependent on cell differentiation. Therefore, PRPF40B may influence transcriptional or post-transcriptional programs independently of TRKB expression, regulating cell cycle progression and coordinating both neuronal proliferation and differentiation. This dual inhibition raises the possibility that PRPF40B functions as a molecular integrator, ensuring the proper balance between proliferation and differentiation. Further investigation into its specific targets and mechanisms could provide insights into how these two processes are interconnected at the molecular level.

In summary, our results highlight the importance of PRPF40B in proliferation, viability, cell migration, and neuronal differentiation by regulating the expression of hundreds of genes, including those involved in neuronal development, such as the TRKB receptor. Further *in vivo* validation studies are needed to fully elucidate the role of PRPF40B in physiological neuronal differentiation and whether PRPF40B is important in pathological conditions associated with impaired BDNF signaling. Such studies will provide valuable information of the potential of PRPF40B as a therapeutic target for neurological disorders.

## METHODS

### Cell lines, Plasmids, and Culture Conditions

SH-SY5Y WT cells were obtained from ATCC. To generate PRPF40B knockout (KO) cell lines using CRISPR/Cas9, two different guide RNAs were employed: gRNA for the G1 cell line (GTAACATCCCCGCGCGGGGG) and gRNA for the G2 cell line (GGCCCGGGCCGTGGCCGATG). The Origene Human Gene Knockout Kit (KN209144) was used according to the manufacturer’s instructions. After selection with puromycin (1.5 µg/mL), individual cell colonies were generated using the limiting dilution method. We also included scrambled control gRNA (Origene, GE100003) to serve as a negative control.

The shRNA target sequence used for SH-SY5Y sh7y(e) was CTAGAGGTTCTAGTCAAACAA (Sigma, TRCN0000075157). Lentiviral production was carried out by transfecting HEK293T cells, cultured in a 10 cm plate, with packaging plasmid, envelope plasmid (1.5 µg each), and shRNA vector DNA (2.5 µg) using the LipoD293 reagent at a 1:3 DNA/reagent ratio, following the manufacturer’s instructions. After 72 hours, the culture medium was harvested, and viral particles were concentrated via ultracentrifugation at 50,000×g for 90 minutes at 4 °C. The virus was then used to infect SH-SY5Y cells, which were subsequently selected with 1.5 µg/mL puromycin. The pLKO.1 plasmid (Sigma) was used as a negative shRNA control.

To generate the MIGR1-GFP-40B construct, wild-type T7-PRPF40B cDNA was subcloned from the pEFBOST7-PRPF40B vector^13^ into EcoRI/BglII-digested MIGR1-GFP vector (Addgene) using PCR. For the retroviral infection, HEK293-T cells were plated in an M6 plate to reach 75-80% confluence after 24 hours. After 24 hours, the medium was aspirated, and 1 mL of DMEM Low medium was added, followed by 1-hour incubation at 37°C. For transfection, a mixture was prepared: Solution A (225 µL of un-supplemented DMEM Low and 15 µL of LipoD [SignaGen Laboratories]) and Solution B (225 µL of un-supplemented DMEM Low, 1.5 µg/µL pVSV-G plasmid, 1.5 µg/µL gag/pol plasmid, and 4 µg/µL MIGR1 empty or MIGR1 PRPF40B plasmid). After the 1-hour incubation, the HEK293-T medium was aspirated, and 400 µL of the A+B solution was added. Cells were incubated for 5 hours at 37°C. Following incubation, the medium was removed, and 5 mL of DMEM medium was added. Cells were incubated at 37°C for 24 hours. For viral particle collection, the retroviral particles were harvested and used to infect the G2 subcellular line. On the same day as the HEK293-T transfection, 400,000 G2 cells were plated in an M6 plate to achieve 80% confluence on the day of infection. 48 hours post-transfection, the medium from HEK293-T cells containing retroviral particles was collected, centrifuged at 448g for 10 minutes, and the supernatant was mixed with polybrene to a final concentration of 4 µg/mL. 2 mL of the mixture was added to the G2 cells, which were then centrifuged at 796 g for 90 minutes, without braking, at 32°C. After centrifugation, unbound virus was removed, cells were added in 2.5 mL of DMEM medium, and were incubated at 37°C for 6 hours. After incubation, the medium was removed and replaced with 2.5 mL of DMEM medium containing puromycin. To assess infection efficiency, the percentage of G2 cells expressing EGFP (from the MIGR1 plasmid) was analyzed by flow cytometry.

SH-SY5Y cell lines were cultured in Dulbecco’s Modified Eagle Medium (DMEM) with high glucose (Invitrogen), supplemented with 10% fetal bovine serum (FBS) and penicillin-streptomycin at final concentrations of 100 U and 100 µg/mL, respectively. Cells were maintained at 37°C in a 5% CO_2_ atmosphere with 95% humidity. SH-SY5Y cells, including SCR, G1, G2, Sh control, Sh7y(e), MIGR1-GFP, and MIGR1-GPF-40B, were grown in the presence of 1 µg/mL puromycin (Invitrogen).

### Cell Proliferation and Colony-Forming Assay

To assess cell proliferation, 2,000 cells per well were seeded into a 96-well plate for each condition. Over a period of 5 days, cells were incubated every 24 hours with 20 µL of Resazurin (100X, Sigma-Aldrich) for 2 hours at 37°C under dark conditions. Fluorescence was measured using an Infinite F200 plate reader (Tecan) at an excitation wavelength of 570 nm and an emission wavelength of 590 nm.

For the colony formation assay, 2,000 cells per well were seeded into a 6-well plate for each condition. Colonies were allowed to grow for one week and then fixed with 1 mL of 4% paraformaldehyde (ChemCruz) for 1 hour at room temperature (RT). After fixation, colonies were stained with 2 mL of 0.1% crystal violet (Sigma) for 1 hour at RT. Stained colonies were dissolved in 2 mL of 20% acetic acid, and the signal intensity was measured using a spectrophotometer (bioNova VersaMax microplate reader) at 590 nm.

### Cell Cycle Analysis

Cells were trypsinized using a 1:3 dilution of 0.05% trypsin-EDTA (Gibco). The cell pellet was washed with 1X PBS and centrifuged at 2,000 rpm for 3 minutes at 4°C. Subsequently, the pellet was resuspended in 100 µL of PBS and 900 µL of 70% ethanol, then incubated for 10 minutes at 4°C. After incubation, the cells were washed again with PBS under the same conditions. The final pellet was resuspended in 300 µL of a solution containing propidium iodide (PI) (40 µg/mL, Sigma) and RNase A (100 µg/mL, Roche). The mixture was incubated for 15 minutes at 37°C in the dark and analyzed using a FACSymphony flow cytometer.

### Apoptosis Assay

Cell pellets were prepared as described for cell cycle analysis. After washing the cell pellet with PBS, the samples were resuspended in 100 µL of Annexin Binding Buffer (10 mM HEPES/NaOH [pH 7.4], 140 mM NaCl, and 2.5 mM CaCl_2_). Control samples not treated with Annexin V were resuspended in 100 µL of PBS. Subsequently, 5 µL of Annexin V-FITC reagent (Immunostep) and/or propidium iodide (PI, 10 µg/mL) was added. The samples were incubated for 15 minutes at RT in the dark. Following incubation, 400 µL of Annexin Binding Buffer or PBS was added, and the samples were analyzed using a FACSymphony flow cytometer. As a control for cell death, 40 µL of 30% H_2_O_2_ (VWR) was added for 15 minutes at 37°C.

### Migration Assay

35 mm micro-Dish plates (Ibidi GmbH) were used according to the manufacturer’s protocol. A total of 200,000 cells were seeded per well, and the insert was removed the following day. Cell migration was monitored by capturing images every 10 minutes over 24 hours using a LEICA DMi8 live-cell microscope.

### Neuronal Differentiation

The protocol described by M.M. Shipley et al.^48^ was followed with modifications. A total of 100,000 cells were seeded per condition (6-well plate or 35 mm dish), and retinoic acid (RA, Sigma) was added the following day. In standard medium (DMEM-high glucose supplemented with 10% FBS and penicillin-streptomycin), RA was added at a final concentration of 10 µM and refreshed every three days. After the RA treatment, cells were washed three times with unsupplemented DMEM-high glucose medium. Subsequently, brain-derived neurotrophic factor (BDNF, 50 ng/µL, CELL guidance systems) was added to the unsupplemented DMEM-high glucose medium, which was also refreshed every three days. Following BDNF treatment, a population of mature neurons was obtained and prepared for further experiments.

### RNA Isolation and RNA-Seq Analysis

For gene expression analysis, RNA was extracted from SH-SY5Y WT, SCR, and G2 cells using the Qiagen RNeasy Micro Kit. RNA concentration and integrity were assessed using a NanoDrop system and an Agilent 2100 Bioanalyzer 2100.

Analysis of sequencing data quality, read distribution (e.g., for potential ribosomal contamination), inner distance size estimation, gene body coverage, and strand specificity of the library were performed using FastQC v0.11.2, Picard-Tools v1.119, Samtools v1.0, and RSeQC v2.3.9. Reads were mapped using STAR v2.7.5a^49^ on the Human hg38 genome assembly, and read count was performed using featureCounts from SubRead v1.5.0 and the Human FAST DB v2021_3 annotations.

Gene expression was estimated as previously described^50^. Only genes expressed in at least one of the two compared conditions were analyzed further. Genes were considered expressed if their FPKM value was greater than FPKM of 98% of the intergenic regions (background). Analysis at the gene level was performed using DESeq2^51^. Genes were considered differentially expressed for fold-changes ≥ 1.5 and p-values ≤ 0.05.

Overrepresented analyses and GSEA^52^ were performed using WebGestalt v0.4.4^53^, merging results from upregulated and downregulated genes, as well as all regulated genes. Pathways and networks were considered significant with p-values ≤ 0.05.

Splicing analysis was first performed by considering only exon reads and flanking exon-exon junction reads (“EXON” analysis) to potentially detect new alternative events that could be differentially regulated (i.e., without considering known alternative events). Splicing analysis was also performed using known splicing patterns (“PATTERN” analysis) based on the FAST DB splicing pattern annotations (i.e., for each gene, all possible splicing patterns are defined by comparing exon content of transcripts). All types of alternative events were analyzed: alternative first exons, alternative terminal exons, cassette exons, mutually exclusive exons, alternative 5’ donor splice sites, alternative 3’ acceptor splice sites, intron retention, internal exon deletion, and complex events corresponding to a mix of several alternative event categories. “EXON” and “PATTERN” analyses are based on the splicing-index calculation as previously described^54–56^. Results were considered statistically significant for p-values ≤ 0.05 and fold-changes ≥ 1.5 in the “PATTERN” analysis, and p-values ≤ 0.01 and fold-changes ≥ 2.0 in the “EXON” analysis. Finally, significant results from “EXON” and “PATTERN” analyses were merged to generate a single result list.

Splice site scores were calculated using MaxEntScan^57^. Polypirimidine tract length, score, and distance of the branch point from the acceptor site were evaluated using SVM-BPfinder^41^.

### RT-qPCR analysis

RNA extraction from all cells was performed using RNA-Solv reagent (Omega Bio-tek). A total of 400 µL of reagent was added to the cell pellet, followed by phenol/chloroform extraction at a 1:1 ratio. The samples were stored at -80°C until further use. Approximately 500 ng of RNA was reverse transcribed using 2 µL of 5X PrimerScript RT Master Mix (Takara Bio Europe) according to the manufacturer’s protocol. To quantify endogenous transcripts, SYBR™ Green PCR Master Mix (Bio-Rad) and the CFX96 Real-Time PCR Detection System were used. Primer sequences are provided in the Supplemental Information (Table S4).

In all RT-qPCR experiments, GAPDH expression was used as an internal control. The quantification of expression was performed using the Pfaffl method^58^, which accounts for primer efficiencies between the gene of interest and the housekeeping gene to improve reproducibility.

### Protein Extraction and Western Blot Analysis

Protein extraction from cell lines was performed using two strong buffers suitable for extracting nuclear and cytoplasmic proteins. For RIPA extraction, the cell pellet was resuspended in 100–500 µL of RIPA buffer (50 mM Tris [pH 7.5], 1% NP-40, 0.05% SDS, 1 mM EDTA, 150 mM NaCl, 0.5% sodium deoxycholate, 1 mM dithiothreitol [DTT], 1 mM phenylmethylsulfonyl fluoride [PMSF], and a protease inhibitor mixture [Complete, Roche]) and incubated for 30 minutes at 4°C. The cell lysate was then centrifuged at maximum speed for 5 minutes at 4°C, and the supernatant was collected. Alternatively, the cells were scraped with 50–500 µL of TR3 buffer (10 mM Na_2_HPO₄, 3% SDS, 10% glycerol, completed with Milli-Q water). The cell extracts were sonicated for 10 seconds at a constant pulse and centrifuged at maximum speed for 5 minutes. The samples were stored at -80°C until use.

Western blotting was performed as previously described^59^. Protein extracts were separated by SDS-PAGE and transferred to a nitrocellulose membrane (Amersham). The primary antibody was incubated overnight (approximately 16 hours) at 4°C. After washing the membrane with PBS-TWEEN, secondary peroxidase-conjugated antibodies were applied for 1 hour at RT. Bound antibodies were detected using ECL™ Start Western Blotting Detection Reagent (Amersham). Antibody sources are provided in the Supplemental Information (Table S5).

### Immunofluorescence

Immunofluorescence in the different SH-SY5Y cell lines was performed as previously described^59^. Images were acquired using the SP8 super-resolution STED confocal microscope. The images were then analyzed and quantified using LAS AF software v2.3.6 and ImageJ (win64). Antibody sources are provided in the Supplemental Experimental Procedures (Table S5).

### Statistics

The statistical analysis was performed using the statistical package included in GraphPad Prism 9.0.1 software. Student’s *t*-test was used to determine significant differences. The data in the graphs represent the mean ± SEM of independent experiments. A p-value < 0.05 was considered significant (*\** p ≤ 0.05, ** p ≤ 0.01, *** p ≤ 0.001, and **** p ≤ 0.0001). Semiquantitative analysis of Western blot bands was performed using Image Lab Software (Bio-Rad). All experiments were performed independently at least three times.

## DATA AVAILABILITY

The accession number for the RNAseq data reported in this paper is GEO: GSE293518. All other data are available in the manuscript or the supplementary materials. Further information and requests for resources and reagents should be directed to and will be fulfilled by the corresponding author.

## ACKNOWLEDGMENTS

The authors would like to thank Juan Valcárcel for critically reading the manuscript and for insightful discussions. The technical assistance of Laura Montosa during the confocal microscopy studies is greatly appreciated. We thank the continued support of Dr. Francisco Javier Oliver (IPBLN) during the final stages of this work.

This work was supported by grants from the Spanish Ministry of Science, Innovation and Universities (grants PID2020-118859GB-I00 to CS and PID2021-128720NB-I00 to C.H.-M.) and the Andalusian Government (grants P20-01269 to C.S. and P20-01271 to C.H.-M.). Support from the European Region Development Fund (ERDF-FEDER) is also acknowledged.

## AUTHOR CONTRIBUTIONS

M.D.-R., A.M.-C., and Y.E.Y designed and performed the experiments, analyzed the data, and revised the manuscript. C.M.-C., N.M.-M., S.J.-L., M.K., C.R.-R., A.R.-C., J.L.-R., and C.H.-M. assisted with the experiments and analyzed data. P.G. performed the bioinformatics analysis. C.H.-M. and C.S. conceptualized and designed the experiments, supervised manuscript preparation, editing and revision, and acquired funding. All authors reviewed and edited the manuscript before submission.

## ADDITIONAL INFORMATION

### Supplementary information

Document S1. Figures S1–S7 and Tables S3 and S4

Table S1. Excel file containing additional data too large to fit in a PDF, related to Figure 3

Table S2. Excel file containing additional data too large to fit in a PDF, related to Figure 4

Table S3. Excel file containing additional data too large to fit in a PDF, related to Figure 4

