## Supplemental Information for "Regulation of *NTRK2* alternative splicing by PRPF40B controls neural differentiation and synaptic plasticity"

### SUPPLEMENTARY INFORMATION

### SUPPLEMENTARY FIGURES

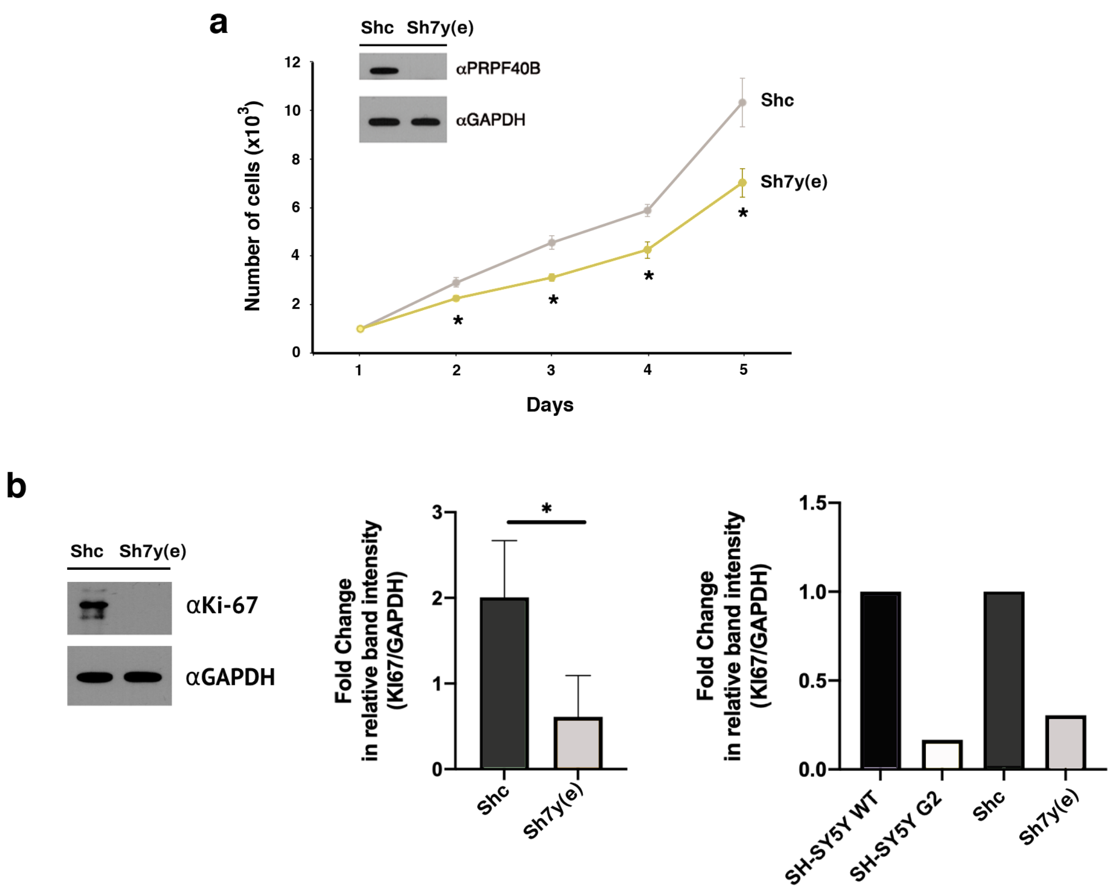


**Supplementary Figure 1. PRPF40B silencing reduces cell viability and proliferation.**

**a** Resazurin assay measuring cell viability in SH-SY5Y cells transfected with either a control shRNA (Shc) aor PRPF40B-specific shRNA (Sh7y(e)) over five consecutive days. Inset: PRPF40B expression was assessed by Western blot using specific antibodies.

**b** Western blot analysis (left) and quantification (right) of Ki-67 expression in Shc and Sh7y(e) cells. Quantification analysis of protein expression using SH-SY5Y WT, G2, Shc, and Sh7y(e) cell lines is also shown.

Data represent the mean ± SEM from three independent experiments. Statistical significance: *$p \leq 0.05$.

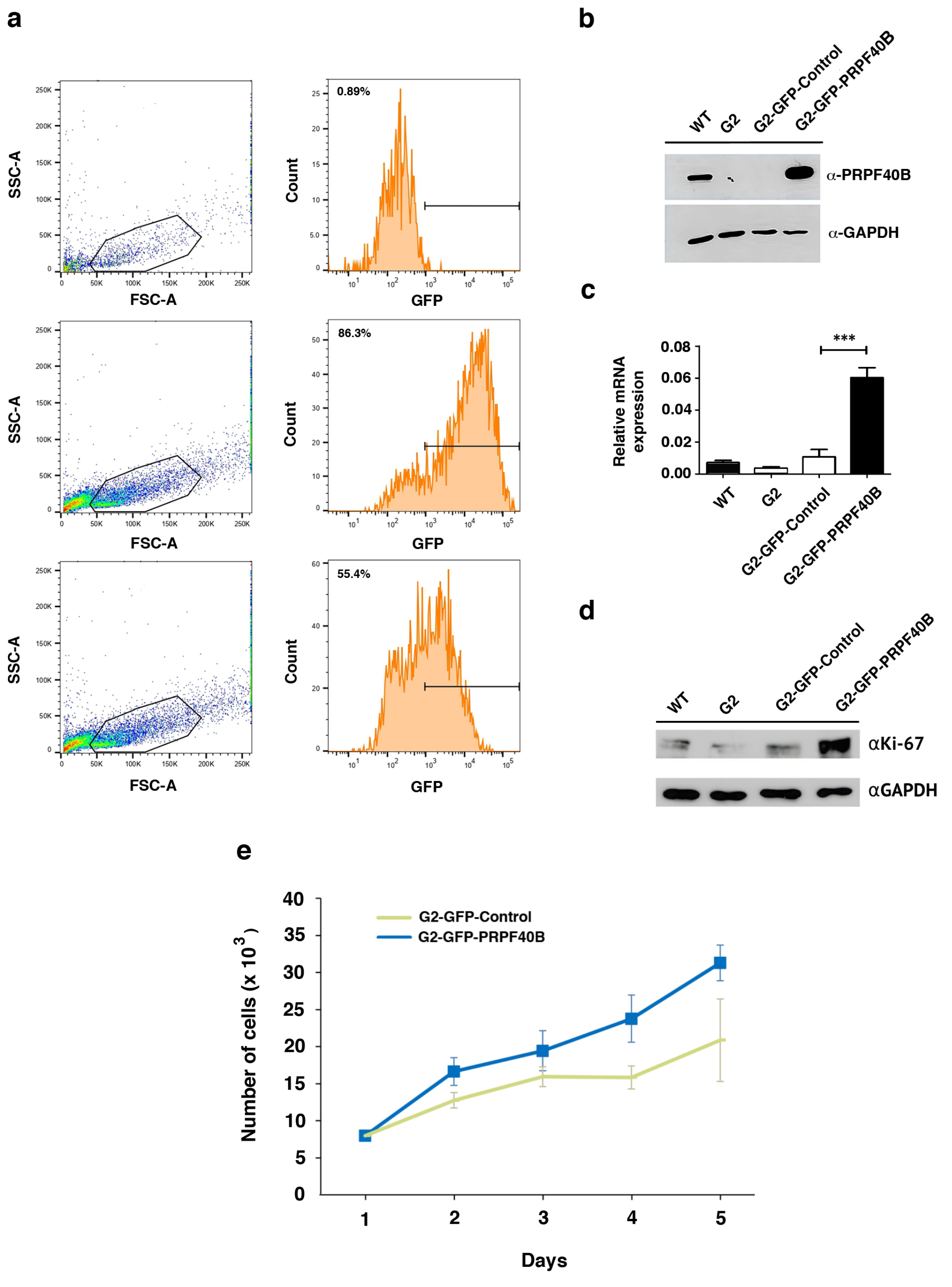


**Supplementary Figure 2. PRPF40B overexpression restores proliferation in G2 cells.**

**a** Graphical representation of flow cytometry results showing GFP expression in G2 cells transduced with either an empty MigR1 retrovirus or retroviruses overexpressing PRPF40B (G2-GFP-Control and G2-GFP-PRPF40B). The transduction efficiency is indicated as a percentage within the graph panels.

**b** Western blot analysis of PRPF40B expression in SH-SY5Y WT, G2, G2-GFP-Control, and G2-GFP-PRPF40B cells. The antibodies used are listed on the right.

**c** Relative mRNA expression levels of PRPF40B in SH-SY5Y WT, G2, G2-GFP-Control, and G2-GFP-PRPF40B cells.

**d** Western blot analysis of Ki-67 expression in SH-SY5Y WT, G2, G2-GFP-Control, and G2-GFP-PRPF40B cells. The antibodies used are listed on the right.

**e** Resazurin assay measuring cell viability in G2-GFP-Control and G2-GFP-PRPF40B cells over five consecutive days.

Data represent the mean ± SEM from three independent experiments. Statistical significance: ***$p \leq 0.001$.

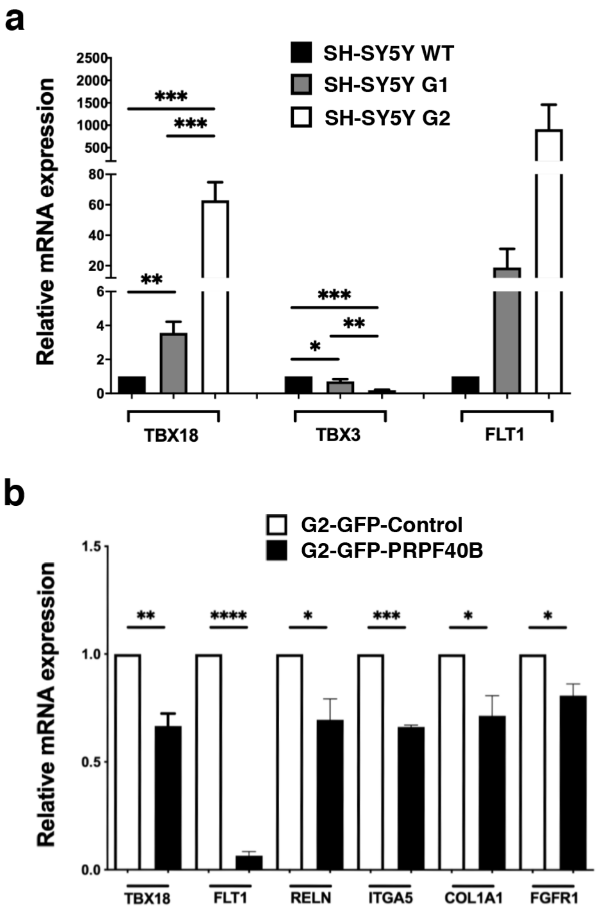


**Supplementary Figure 3. RT-qPCR validation of differentially expressed genes.**

**a** RT-qPCR analysis of selected DE genes in SH-SY5Y WT, G1, and G2 cells.

**b** RT-qPCR analysis of the indicated DE genes in G2-GFP-Control and G2-GFP-PRPF40B cells.

Data represent the mean ± SEM from three to six independent experiments. Statistical significance: $^{*}p \leq 0.05$, $^{**}p \leq 0.01$, $^{***}p \leq 0.001$, and $^{****}p \leq 0.0001$.

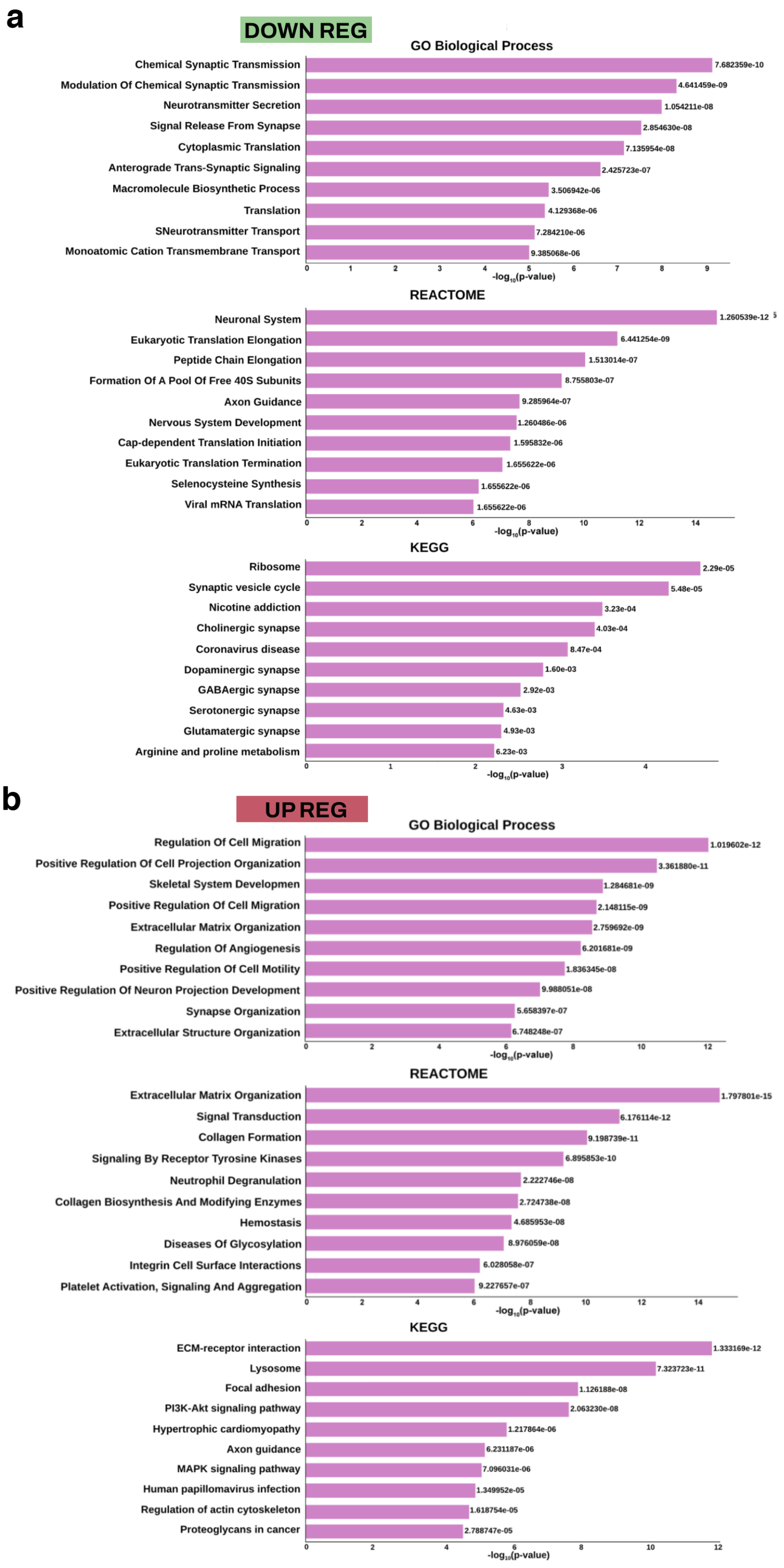
a
DOWN REG
GO Biological Process
Chemical Synaptic Transmission
7.682359e-10
Modulation Of Chemical Synaptic Transmission
4.641459e-09
Neurotransmitter Secretion
1.054211e-08
Signal Release From Synapse
2.854630e-08
Cytoplasmic Translation
7.135954e-08
Anterograde Trans-Synaptic Signaling
2.425723e-07
Macromolecule Biosynthetic Process
3.506942e-06
Translation
4.129368e-06
SNeurotransmitter Transport
7.284210e-06
Monoatomic Cation Transmembrane Transport
9.385068e-06
-log10(p-value)
REACTOME
Neuronal System
1.260539e-12
Eukaryotic Translation Elongation
6.441254e-09
Peptide Chain Elongation
1.513014e-07
Formation Of A Pool Of Free 40S Subunits
8.755803e-07
Axon Guidance
9.285964e-07
Nervous System Development
1.260486e-06
Cap-dependent Translation Initiation
1.595832e-06
Eukaryotic Translation Termination
1.655622e-06
Selenocysteine Synthesis
1.655622e-06
Viral mRNA Translation
1.655622e-06
-log10(p-value)
KEGG
Ribosome
2.29e-05
Synaptic vesicle cycle
5.48e-05
Nicotine addiction
3.23e-04
Cholinergic synapse
4.03e-04
Coronavirus disease
8.47e-04
Dopaminergic synapse
1.60e-03
GABAergic synapse
2.92e-03
Serotonergic synapse
4.63e-03
Glutamatergic synapse
4.93e-03
Arginine and proline metabolism
6.23e-03
-log10(p-value)
b
UP REG
GO Biological Process
Regulation Of Cell Migration
1.019602e-12
Positive Regulation Of Cell Projection Organization
3.361880e-11
Skeletal System Developmen
1.284681e-09
Positive Regulation Of Cell Migration
2.148115e-09
Extracellular Matrix Organization
2.759692e-09
Regulation Of Angiogenesis
6.201681e-09
Positive Regulation Of Cell Motility
1.836345e-08
Positive Regulation Of Neuron Projection Development
9.988051e-08
Synapse Organization
5.658397e-07
Extracellular Structure Organization
6.748248e-07
-log10(p-value)
REACTOME
Extracellular Matrix Organization
1.797801e-15
Signal Transduction
6.176114e-12
Collagen Formation
9.198739e-11
Signaling By Receptor Tyrosine Kinases
6.895853e-10
Neutrophil Degranulation
2.222746e-08
Collagen Biosynthesis And Modifying Enzymes
2.724738e-08
Hemostasis
4.685953e-08
Diseases Of Glycosylation
8.976059e-08
Integrin Cell Surface Interactions
6.028058e-07
Platelet Activation, Signaling And Aggregation
9.227657e-07
-log10(p-value)
KEGG
ECM-receptor interaction
1.333169e-12
Lysosome
7.323723e-11
Focal adhesion
1.126188e-08
PI3K-Akt signaling pathway
2.063230e-08
Hypertrophic cardiomyopathy
1.217864e-06
Axon guidance
6.231187e-06
MAPK signaling pathway
7.096031e-06
Human papillomavirus infection
1.349952e-05
Regulation of actin cytoskeleton
1.618754e-05
Proteoglycans in cancer
2.788747e-05
-log10(p-value)

**Supplementary Figure 4. GO, REACTOME, and KEGG pathway enrichment analyses of differentially expressed genes.**

GO term enrichment analysis (Biological process domain), along with REACTOME and KEGG pathway enrichment analyses, was performed using all differentially downregulated (**a**) and upregulated (**b**) genes detected upon PRPF40B silencing. A subset of the top enriched pathways and their –log10 enrichment values are shown.

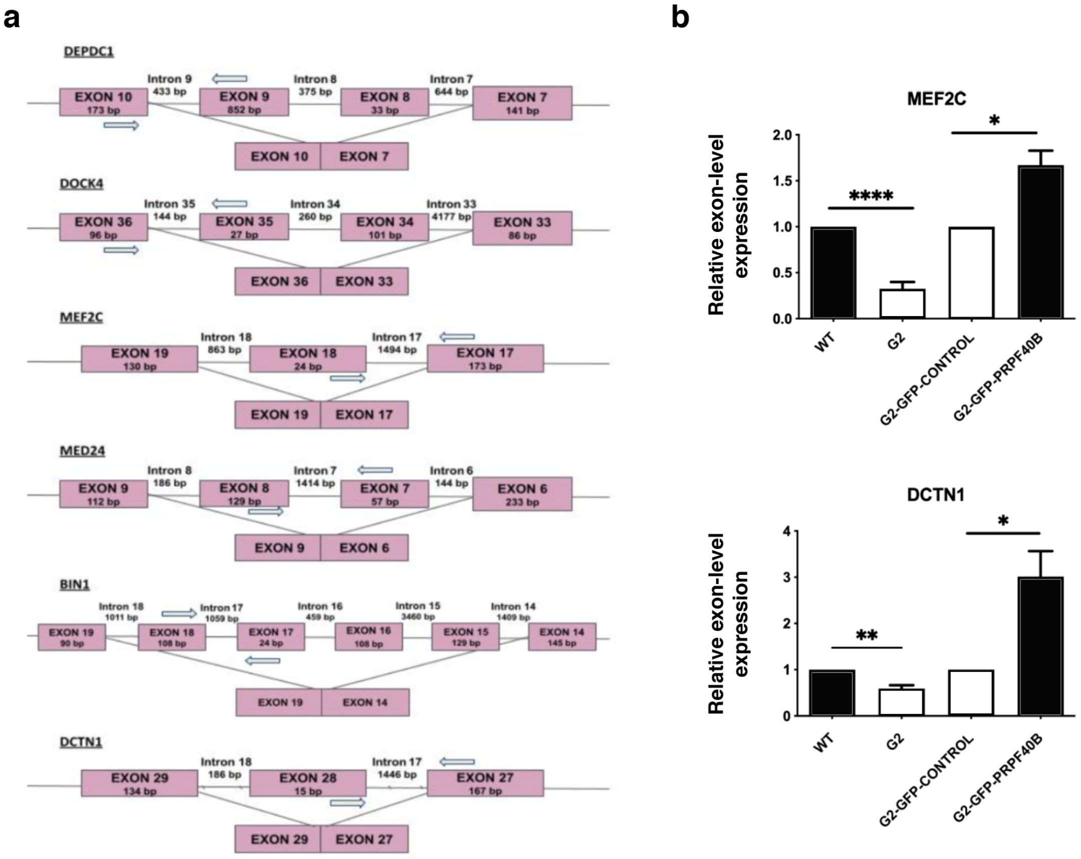


**Supplementary Figure 5. Analyses of RNA-seq data for alternative splicing events.**

**a** Schematic representation of each validated target gene (Figure 4E), showing exons (boxes), introns (lines), and the predicted alternative processing event. Arrows indicate primer locations.

**b** RT-qPCR analysis of the indicated genes in SH-SY5Y WT, G2, G2-GFP-Control, and G2-GFP-PRPF40B cells, demonstrating the restoration of regulated exon expression upon PRPF40B overexpression.

Data represent the mean ± SEM from three independent experiments. Statistical significance: $*p \leq 0.05$, $**p \leq 0.01$, and $****p \leq 0.0001$.

**a**

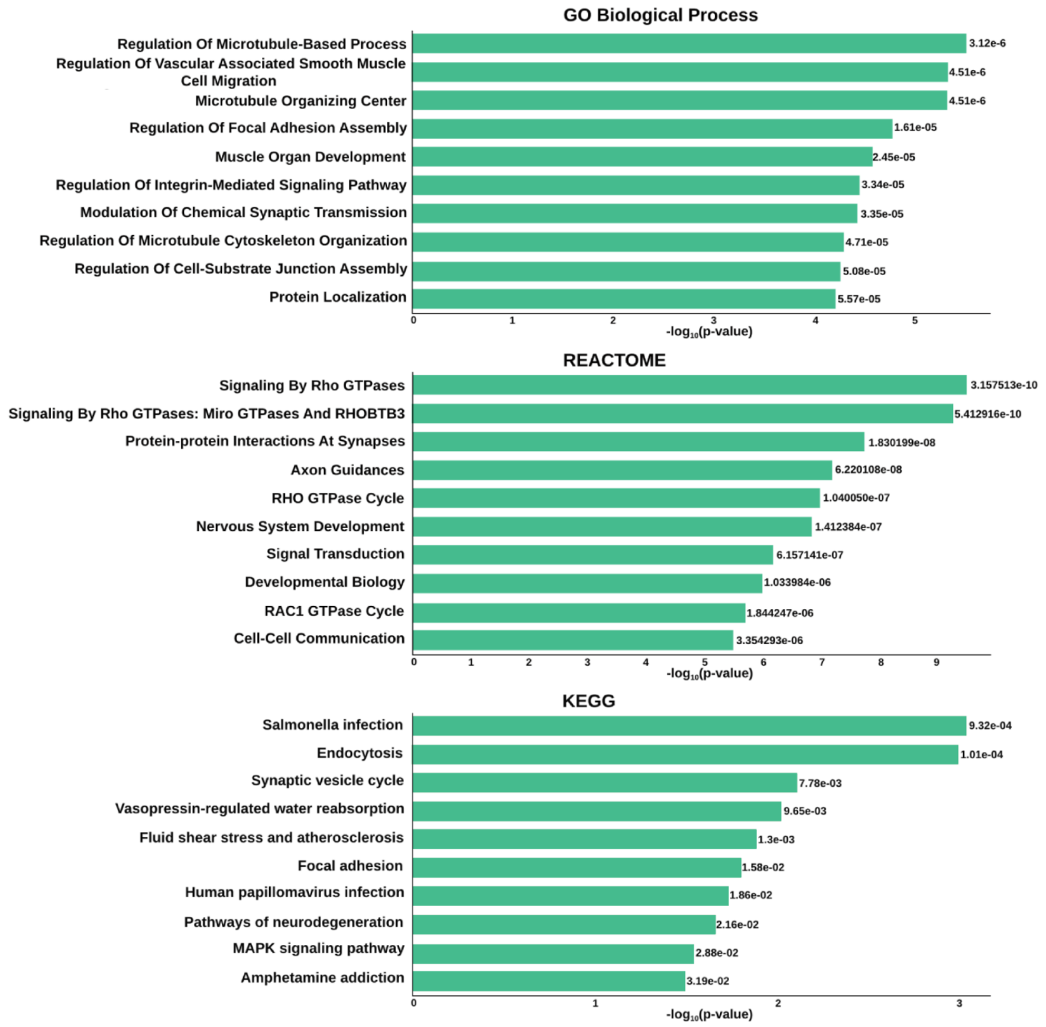


**b**

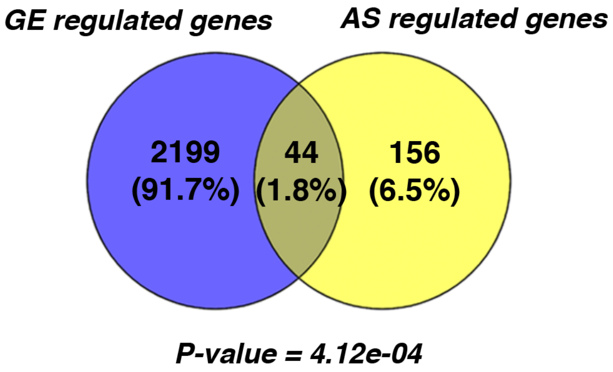


**Supplementary Figure 6. GO, REACTOME, and KEGG pathway enrichment analyses of differentially expressed splicing events.**

**a** GO term enrichment analysis (Biological process domain), along with REACTOME and KEGG pathway enrichment analyses, was performed using all differentially regulated alternative splicing events.

**b** Overlap of genes with differential gene expression and differential alternative splicing.

**a**

**b**

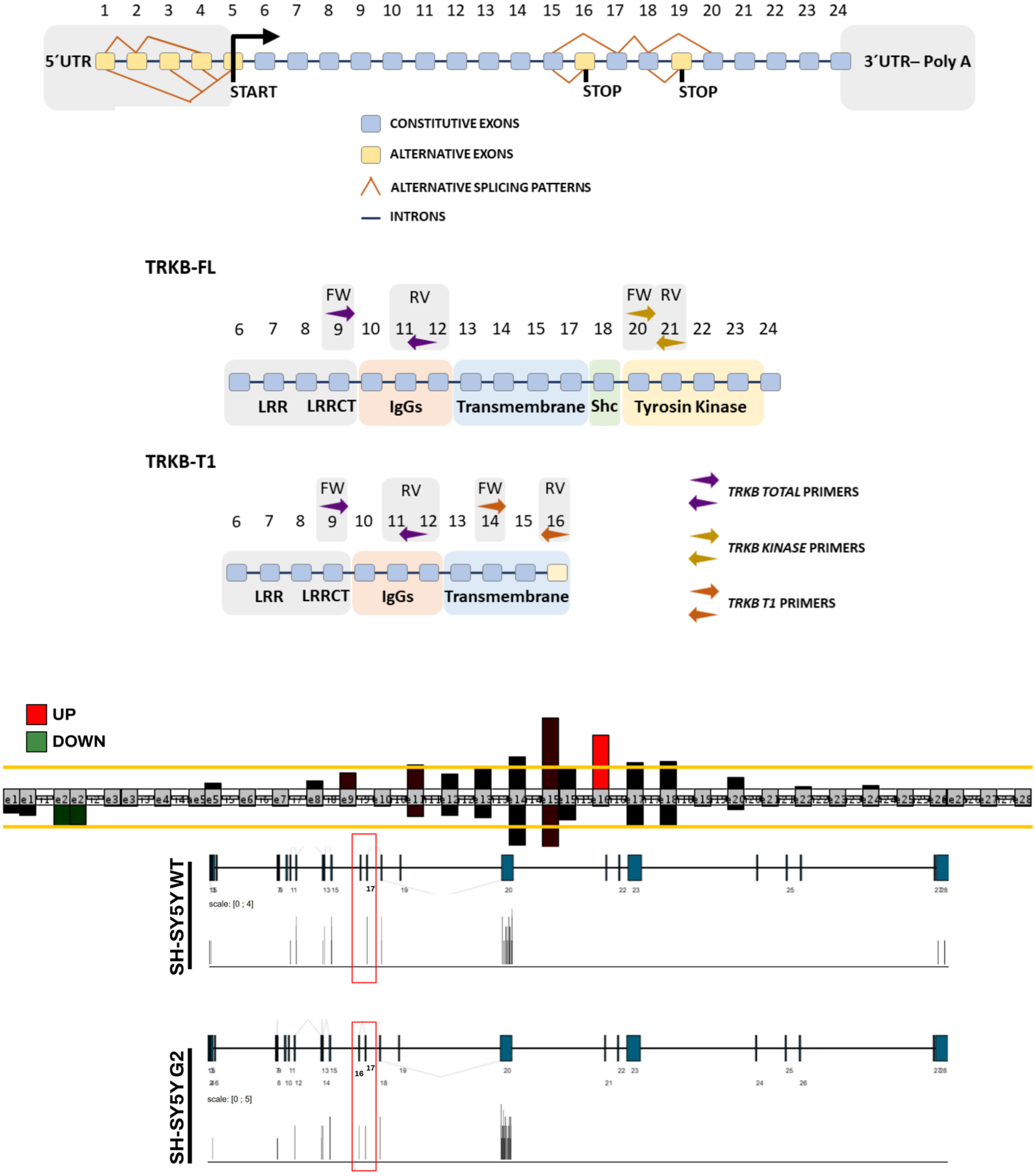


**Supplementary Figure 7. Alternative splicing of *NTRK2* and its regulation by PRPF40B.**

**a** Schematic representation of *NTRK2* pre-mRNA. Exons and introns are depicted as boxes and horizontal lines, respectively. Alternative splicing patterns are represented by red lines connecting exons. Constitutive exons are shown in blue, while alternative exons are shown in yellow. The positions of two alternative stop codons are also indicated. Below, the two main mature transcripts derived from *NTRK2* pre-mRNA (TRKB-FL and TRKB-T1) through alternative splicing are illustrated. The predicted domains of TRKB, including Leucine-rich repeats (LRR), Leucine-rich repeat C-terminal domain (LRRCT), Immunoglobulin-like domains (IgGs), transmembrane domain, Src homology 2 domain (Shc), and the protein kinase domain, are shown. The positions of primers used to detect total TRKB, TRKB-FL, and TRKB-T1 are indicated above the diagrams.

**b** Representation of the *NRK2* gene cluster using the EASANA visualization module. Exon and their corresponding numbers are shown in gray rectangles. Each bar represents a probe. Bar height indicates the mean signal intensity (log 2 scale) under control or PRPF40B-depleted conditions. Red bars represent upegulated probes, green bars indicate downregulated probes, and black bars correspond to probes with no intensity change between conditions. Below the diagram, a magnified schematic highlights exon 16 change.

**SUPPLEMENTARY TABLES**

**Supplementary Table 4: List of Oligonucleotides**

| NAME | SEQUENCE |
|---|---|
| ***TRANSCRIPTION*** | |
| SPARC FW | 5´-ATGGTGCAGAGGAAACCGAA-3´ |
| SPARC RV | 5´-TTCTCATCCAGCTCGCACAC-3´ |
| GAP43 FW | 5´-CCATGCTGTGCTGTATGAGAAGAAC-3´ |
| GAP43 RV | 5´-AAGCAAGGGCTGAGGTGTTA-3´ |
| TBX18 FW | 5´- CTTCGACAAGCTGAAGCTCAC-3´ |
| TBX18 RV | 5´- GGCTTGATGGGAGAAAGATCG-3´ |
| TBX3 FW | 5´- TCCCAAGTGATCACGCTACG-3´ |
| TBX3 RV | 5´-GTGTCCCGGAAACCTTTTGC-3´ |
| IGFBP3 FW | 5´- GACTACGAGTCTCAGAGCAC-3´ |
| IGFBP3 RV | 5´- TCTTGTCACAGTTGGGAATGTG-3´ |
| RELN FW | 5´- TCGTCCTAGTAAGCACTCGCA-3´ |
| RELN RV | 5´- ATCGCCTAAGTGACCTTCGT-3´ |
| ITGA5 FW | 5´- GGCTTCAACTTAGACGCGGA-3´ |
| ITGA5 RV | 5´- GGCCGGTAAAACTCCACTGA-3´ |
| COL1A1 FW | 5´- TAGGGTCTAGACATGTTCAGCTTTGT-3´ |
| COL1A1 RV | 5´- GTGATTGGTGGGATGTCTTCGT-3´ |
| FGFR1 FW | 5´- TCAGATGCTCTCCCCTCTC-3´ |
| FGFR1 RV | 5´- TACGGGCATACGGTTTGGTT-3´ |
| GAPDH FW | 5´- TACGGGCATACGGTTTGGTT-3´ |
| GAPDH RV | 5´- GGGTCATTGATGGCAACAATATC-3´ |
| FLT1 FW | 5´-CGGTCTTACCGGCTCTCTAT-3´ |
| FLT1 RV | 5´-GCCACGAGTCAAATAGCGAG-3´ |
| NTRK2.Total FW | 5´- CTGGCCTGGAATTGACGATG-3´ |
| NTRK2.Total RV | 5´- ACCACAGCATAGACCGAGAG-3´ |
| ***SPLICING*** | |
| NTRK2.KINASE FW | 5´- GGGTCGTGTTGTGGGAGATT-3´ |
| NTRK2.KINASE RV | 5´- TGTTCTTCCTCATGTGGGGC-3´ |
| NTRK2.T1 FW | 5´- GGGGACACCACGAACAGAAG-3´ |
| NTRK2.T1 RV | 5´- CTACCCATCCAGTGGGATCTT-3´ |
| DEPDC1 FW | 5’- TCGGAGAGTCTAGTGCCACA -3’ |
| DEPDC1 RV | 5’- ACTGTAGAGCATCGATGGCA -3’ |
| DOCK4 FW | 5’- TCCACTATTTGGTCCCTATCCT -3’ |
| DOCK4 RV | 5’- AACAACCTCTCCATTAGACGAG -3’ |

| | |
|---|---|
| MEF2C FW | 5'- GCAAGCAAAATCTCCTCCCC -3' |
| MEF2C RV | 5'- AAAGCAGGTCGACATCCTCA -3' |
| MED24 FW | 5'- CCTGGAGAAAACCCTCAGCA -3' |
| MED24 RV | 5'- CTGTTGGTTGATTGGCTCGG -3' |
| BIN1 FW | 5'- ATTCCATGGGACCTCTGGGA -3' |
| BIN1 RV | 5'- GCCAGGACACAGCAAAGGT -3' |
| DCTN1 FW | 5'- AGGCAGAACTAAAGCAGCGT -3' |
| DCTN1 RV | 5'- TGTTCTCATGCTGGAGCTGG -3' |

**Supplementary Table 5: List of Antibodies**

| NAME | TECHNIQUE | REFERENCE | DILUTION |
|---|---|---|---|
| αPRPF40B | IF | Proteintech/16929-1-AP | 1:100 |
| αPRPF40B | WB | Abcam/ab122474 | 1:250 |
| αKI-67 | WB | BD Pharmingen/550609 | 1:500 |
| αFAK$^{Y397}$ | IF | Abcam/ab81298 | 1:1000 |
| αβ-Tubulin III | WB | Sigma-Aldrich/T8660 | 1:2000 |
| αβ-Tubulin III | IF | Sigma-Aldrich/T8660 | 1:200 |
| αPI3K | WB | Santacruz/sc-365290 | 1:250 |
| αAKT | WB | CellSignaling/#92725S | 1:500 |
| αp-AKT | WB | CellSignaling/#9271 | 1:500 |
| αERK 1/2 | WB | CellSignaling/#4695 | 1:500 |
| αp-ERK 1/2 | WB | CellSignaling/#4370 | 1:1000 |
| αDrebrin | WB | Santacruz/sc-374269 | 1:100 |
| αSYP | WB | Santacruz/sc-17750 | 1:500 |
| αSynapsin 1a/b | WB | Santacruz/sc-376623 | 1:100 |
| αComplexin 1/2 | WB | Santacruz/sc-365152 | 1:250 |
| αTRKB | WB | Proteintech/13129-1-AP | 1:100 |
| αN-Cadherin | WB | Santacruz/sc-393933 | 1:500 |
| αGAPDH | WB | Santacruz/sc-365062 | 1:5000 |
| αIgGRabbit-HRP | WB | Abcam/ab205718 | 1:5000 |
| αIgGMouse-HRP | WB | Dako/P0447 | 1:5000 |
| αIgGMouse-Alexa488 | IF | Molecular Probes/A-32723 | 1:500 |
| αIgGRabbit-Alexa647 | IF | Molecular Probes/A-21245 | 1:500 |
